# The biogeography of *Streptomyces* in New Zealand enabled by high-throughput sequencing of genus-specific *rpoB* amplicons

**DOI:** 10.1101/2020.09.22.309195

**Authors:** S.A. Higgins, K. Panke-Buisse, Daniel H. Buckley

## Abstract

We evaluated *Streptomyces* biogeography in soils along a 1,200 km latitudinal transect across New Zealand (NZ). *Streptomyces* diversity was examined using high-throughput sequencing of *rpoB* amplicons generated with a *Streptomyces* specific primer set. We detected 1,287 *Streptomyces rpoB* operational taxonomic units (OTUs) with 159 ± 92 (average ± s.d.) *rpoB* OTUs per site. Only 12% (n = 149) of these OTUs matched *rpoB* sequences from cultured specimens (99% nucleotide identity cutoff). *Streptomyces* phylogenetic diversity (Faith’s PD) was correlated with soil pH, mean annual temperature, and plant community richness (Spearman’s *r*: 0.77, 0.64, and −0.79, respectively; p < 0.05), but not with latitude. In addition, soil pH and plant community richness both explained significant variation in *Streptomyces* beta diversity. *Streptomyces* communities exhibited both high dissimilarity and strong dominance of one or a few species at each site. Taken together, these results suggest that dispersal limitation due to competitive interactions limits the colonization success of spores that relocate to new sites. Cultivated *Streptomyces* isolates represent a major source of clinically useful antibiotics, but only a small fraction of extant diversity within the genus have been identified and most species of *Streptomyces* have yet to be described.

## Introduction

The genus *Streptomyces* consists of a cosmopolitan group of spore-forming microorganisms whose ease of cultivation has facilitated their investigation for over a century (Waksman *et al*., 1948; Kämpfer *et al*., 2014). They are aerobic, filamentous members of the gram-positive bacterial phylum *Actinobacteria*, possessing characteristic developmental, biosynthetic, and catabolic genes which have contributed to their success in a diversity of terrestrial and aquatic habitats for approximately 350 myr (Seipke *et al*., 2012; Kämpfer *et al*., 2014; McDonald and Curriea, 2017). Furthermore, their utility as model organisms in investigations of natural product synthesis, cell biology, and evolution has transformed microbiology and biotechnology research (Kämpfer *et al*., 2014; Chater, 2016). While their capacity for carbon mineralization (Ruddick and Williams, 1972; Barder and Crawford, 1981; Spiker *et al*., 1992; Dari *et al*., 1995; Schlatter *et al*., 2009) and secondary metabolism (Chater, 2016; Niu *et al*., 2016; Gallagher *et al*., 2017; Choudoir *et al*., 2018) have been well-documented, much less is known regarding their diversity, ecology, and biogeography in terrestrial habitats. To date, biogeographical studies of the *Streptomyces* have largely focused on genetic variation within cultivated representatives to inform population level analyses (Davelos *et al*., 2004; Antony-Babu *et al*., 2008; Andam *et al*., 2016). While instructive, these investigations are limited to the cultivable minority of *Streptomyces* species and are likely insufficient to capture the breadth of localized, ecological processes shaping *Streptomyces* biodiversity and community structure *in situ*. Nevertheless, cultivation-based approaches have enabled valuable physiological investigations of diverse traits within members of the *Streptomyces* (Schlatter *et al*., 2013; Choudoir and Buckley, 2018), yet cultivation-independent explorations of the structure of *Streptomyces* communities are lacking and represent an underexplored area of research regarding the ecology of this diverse genus.

The universality of PCR primers targeting the 16S rRNA (16S) gene across bacterial taxa has facilitated key scientific investigations related to expanding our understanding of microbial diversity (Baker *et al*., 2003; Lauber *et al*., 2009). However, the significant conservation within the 16S gene is indicative of a slow evolutionary rate (1% nucleotide divergence ∝ 50 myr) permitting coarse diversity profiling, but is insufficient to detect fine-grained population structure over recent evolutionary time scales (i.e., tens to hundreds of thousands of years) (Choudoir *et al*.; Ochman and Wilson, 1987; Ochman *et al*., 1999; Adékambi *et al*., 2009). For diversity assessments within some congeneric microbial lineages, the 16S gene is inadequate to delineate “species” or fine-grain operational taxonomic units (OTUs), and alternative genes have been identified that improve discrimination where fine scale diversity trends are of interest (Case *et al*., 2007; Antony-Babu *et al*., 2017). The gene encoding the beta subunit of the DNA-directed RNA polymerase (*rpoB*) has been shown to differentiate closely-related *Streptomyces* species when the 16S gene could not (Kim *et al*., 2004; Case *et al*., 2007). Hence, protein-coding genes, such as *rpoB*, are preferred for biogeographic explorations of microorganisms and may facilitate identification of underlying community assembly mechanisms (Koeppel and Wu, 2013).

While cultivation-dependent investigations of *Streptomyces* species have uncovered novel biogeographic patterns related to *Streptomyces* species diversity (Choudoir *et al*., 2016; Choudoir and Buckley, 2018), these isolation-based studies are laborious and do not scale to the large number of samples required for investigations of broad-scale biogeographic phenomena. Hence, we set out to develop and apply *Streptomyces*-specific *rpoB* PCR primers compatible with high-throughput Illumina sequencing to investigate *Streptomyces* community structure in environmental samples. To demonstrate the utility of the *Streptomyces rpoB* primers, we amplified and sequenced *Streptomyces rpoB* genes from soils spanning 1,200 km along a latitudinal transect in New Zealand (NZ). Biogeographical studies in NZ have uncovered distinctive patterns of floral and faunal diversity trends owing to the continental island’s unique paleoclimatic and geologic history (Fleming, 1975; Heads, 1997; Wallis and Trewick, 2009; McGlone *et al*., 2010; Tonkin *et al*., 2018), and information regarding the diversity and structure of *Streptomyces* communities across this temperate island region are lacking. Historical phenomena (e.g., glaciation, climatic anomalies) are correlated with both *Streptomyces* phylogenetic diversity and total bacterial community richness (Eisenlord *et al*., 2012; Choudoir *et al*., 2016; Delgado-Baquerizo *et al*., 2017), but the degree to which paleo and contemporary climate shape *Streptomyces* diversity across NZ is unknown and may advance our understanding of bacterial diversity trends within NZ soils. Therefore, we performed high-throughput sequencing of *Streptomyces rpoB* amplicons along a latitudinal transect in NZ to investigate ecological processes (selection, drift, dispersal, diversification) shaping *Streptomyces* communities in soil. Furthermore, we test the hypothesis that phylogenetic clustering of *Streptomyces* communities will occur poleward, consistent with a legacy of widespread glaciation on NZ’s south island as has previously been observed in North America (Andam *et al*., 2016).

## Results and Discussion

### Determining Streptomyces diversity by rpoB analysis

#### Patterns of rpoB alpha diversity

Using *Streptomyces* specific *rpoB* primers we observed 159 ± 92 *Streptomyces rpoB* OTUs per site (mean ± s.d., range: 53 to 395) across the 1,200 km NZ latitudinal transect. When using 16S gene data to identify *Streptomyces* OTUs, only 9 ± 2 OTUs and 16 ± 5 ESVs were observed per site. Accordingly, the *rpoB* primers demonstrated higher resolution than 16S gene primers for detecting diversity within the genus S*treptomyces* (Fig. S1). This finding corroborates previous observations that the 16S gene lacks sufficient genetic resolution to discriminate between *Streptomyces* species (Labeda et al. 2017, Cheng et al. 2016, Rong et al. 2013), and that *rpoB* is a preferred taxonomic marker for members of the *Streptomyces* (Andam 2016). The rate of new species discovery using *rpoB* sequences declined rapidly as read counts approached 1,000 (Fig. S2). At a sequence depth of 5,000 counts per sample, *rpoB* OTU coverage met or exceeded 99% for all but two samples (Table S1). Sequencing coverage for *rpoB* OTUs in each site ranged between 0.95 and 0.99 at a read count of 922 sequences (Table S1), the lowest read count observed in this study. Thus, amplicon pools generated with the *Streptomyces rpoB* primers can be multiplexed to determine near complete *Streptomyces* species inventories across large numbers of soil samples.

Of the four abiotic variables examined, pH and MAT were significantly correlated with *Streptomyces* phylogenetic diversity (PD) (Fig. 1A, Table 1), whereas MAR and MAWS were non-significant and negatively correlated with PD (Fig. 1B, Table 1). No significant correlation was detected between *Streptomyces* PD and elevation. In addition to abiotic variables, we observed a significant negative correlation between *Streptomyces* PD and NZ plant species richness (Table 1). While clear links exist between *Streptomyces* community structure and plant growth promotion or plant host genotype (Seipke *et al*., 2012; Bakker *et al*., 2013; Viaene *et al*., 2016; Essarioui *et al*., 2017), additional experiments are needed to support a causal influence of plant diversity on *Streptomyces* diversity. Aside from their beneficial effects on plants, a minority of *Streptomyces* species also represent harmful plant pathogens that cause scabs lesions on root storage organs (e.g., potato) (Loria *et al*., 2006; Anderson *et al*., 2014). Of the 1,287 *rpoB* OTUs, only 10 OTUs matched *rpoB* sequences classified as canonical scab-causing *Streptomyces* species. However, these OTUs were detected in 6 out of 15 NZ soil samples, and ranged in abundance from 0 to 3.8% (0.5 ± 1.1 %; mean ± s.d.) of sequences in a given sample. Although additional evidence is required to confirm the presence of pathogenic *Streptomyces* species in the NZ soils, our results suggest that high-throughput *rpoB* sequencing could provide an alternative, cultivation-independent method to surveil putatively pathogenic populations of *Streptomyces* spp. in soils.

**Table 1.** Spearman rank correlations between *Streptomyces* diversity and select variables estimated for the NZ soil samples (*n* = 15). Diversity measurements reported are log_10_ of observed *rpoB* OTUs (OTU_obs_), Faith’s phylogenetic diversity (PD), community evenness (E_var_), and net relatedness index (NRI). Significant results (*p* < 0.05) are indicated by bold font.

|  | pH | MAR | MAT | MAWS | Latitude | Plant richness |
| --- | --- | --- | --- | --- | --- | --- |
| $\text{OTU}_{\text{obs}}$ | 0.42 | **-0.59** | 0.08 | -0.41 | -0.08 | -0.44 |
| PD | **0.77** | -0.32 | **0.64** | -0.03 | 0.40 | **-0.79** |
| $E_{\text{var}}$ | 0.38 | **-0.67** | 0.15 | -0.28 | -0.25 | -0.34 |
| NRI | 0.30 | -0.15 | 0.32 | 0.11 | 0.30 | **-0.73** |

MAR = mean annual rainfall (mm), MAT = mean annual temperature ( $^{\circ}\text{C}$ ), MAWS = mean annual wind speed (m/s).

**Figure 1.**
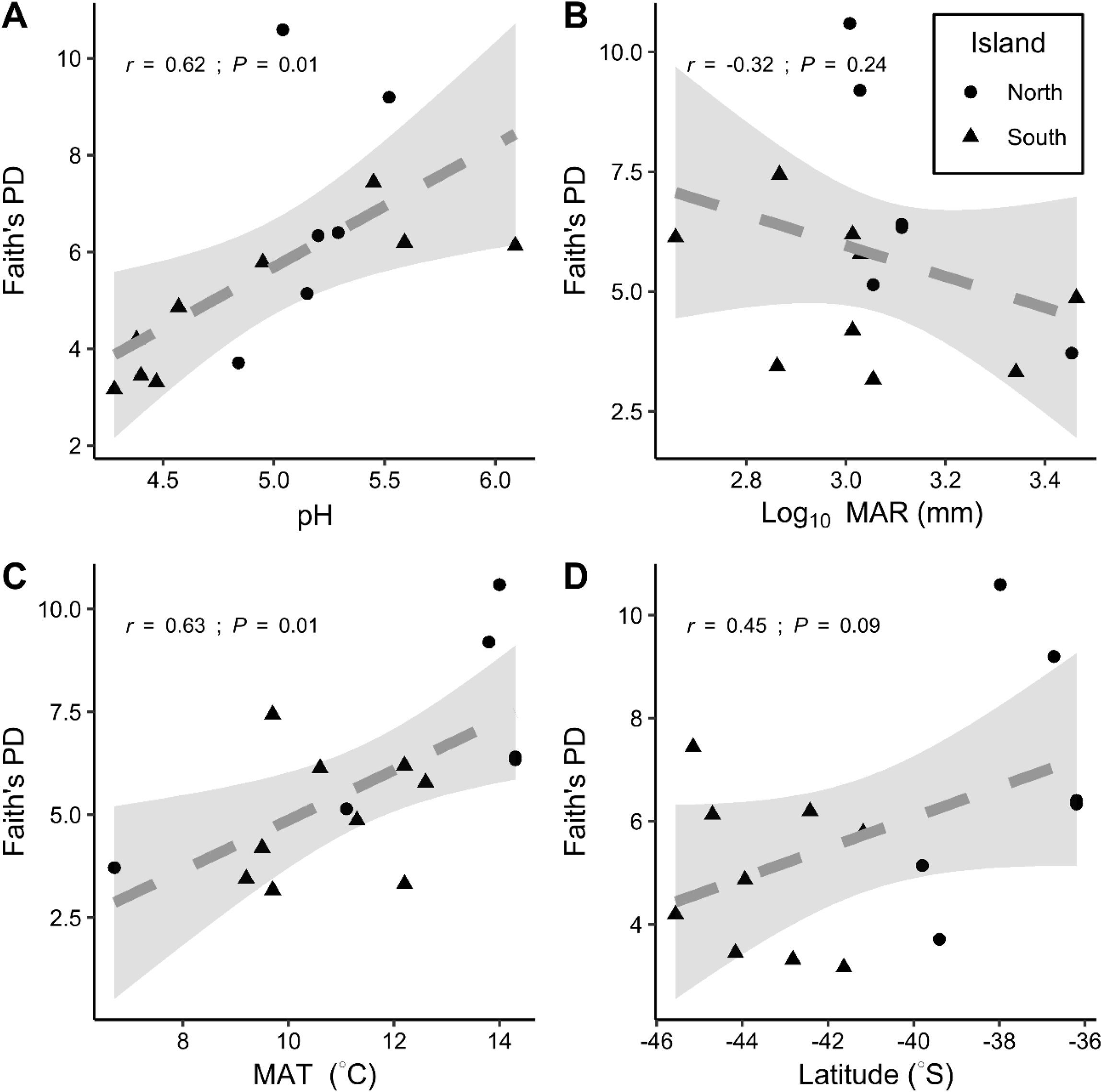
Association between *Streptomyces* phylogenetic diversity (PD) and environmental parameters. MAR = mean annual rainfall; MAT = mean annual temperature; *r* = spearman rank correlation.

#### Patterns of rpoB evenness

Estimates of *Streptomyces* community evenness were low (0.19 ± 0.06, mean ± s.d.) and only MAR demonstrated a significant negative correlation with E_var_ (Table 1). The dominant *rpoB* OTU comprised up to 60% (31 ± 15 %, mean ± s.d.) of *Streptomyces* sequences observed within each site, indicating strong dominance within *Streptomyces* communities in NZ soils. *Streptomyces* spp. are ubiquitous and easily cultivated (Kämpfer *et al*., 2014), but additional studies are required to identify if a cultivation bias exists that selects for numerically dominant *Streptomyces* spp. in soils. Although changes in community composition during storage could explain the strong dominance we observed, electrophoresis analysis of *Streptomyces* 16S gene profiles from fresh soils also display dominance (Inbar *et al*., 2005). In addition, when we compared soils in long term dry storage to those frozen rapidly we did not observe a systematic bias towards dominance (Fig. S3).

### Streptomyces beta diversity based on rpoB

#### Spatial, historical, and climatic correlates of Streptomyces beta diversity

*Streptomyces* community composition was highly dissimilar between sites and did not correlate with geographic distance (Fig. 2; Mantel r < 0.05, *P* > 0.4; Procrustes Gower statistic, m_12_ = 0.81 – 0.86, *P* > 0.36). Furthermore, no difference in *Streptomyces* community composition was detected between NZ’s north and south islands when assessed using either *rpoB* (F_1,13_ = 1.17, R^2^ = 0.08, *p* > 0.1) or 16S gene analyses (F_1,13_ = 1.15, R^2^ = 0.08, *p* > 0.2) (Fig. 3). Instead, soil pH was correlated with both total bacterial and *Streptomyces* community composition across NZ (Fig. 3), as previously observed for soil bacterial communities (Lauber *et al*., 2009). Hence, habitat filtering via pH regimes might play a strong role in structuring soil *Streptomyces* communities as has been observed generally for soil *Actinobacteria* (Lauber *et al*., 2009).

**Figure 2.**
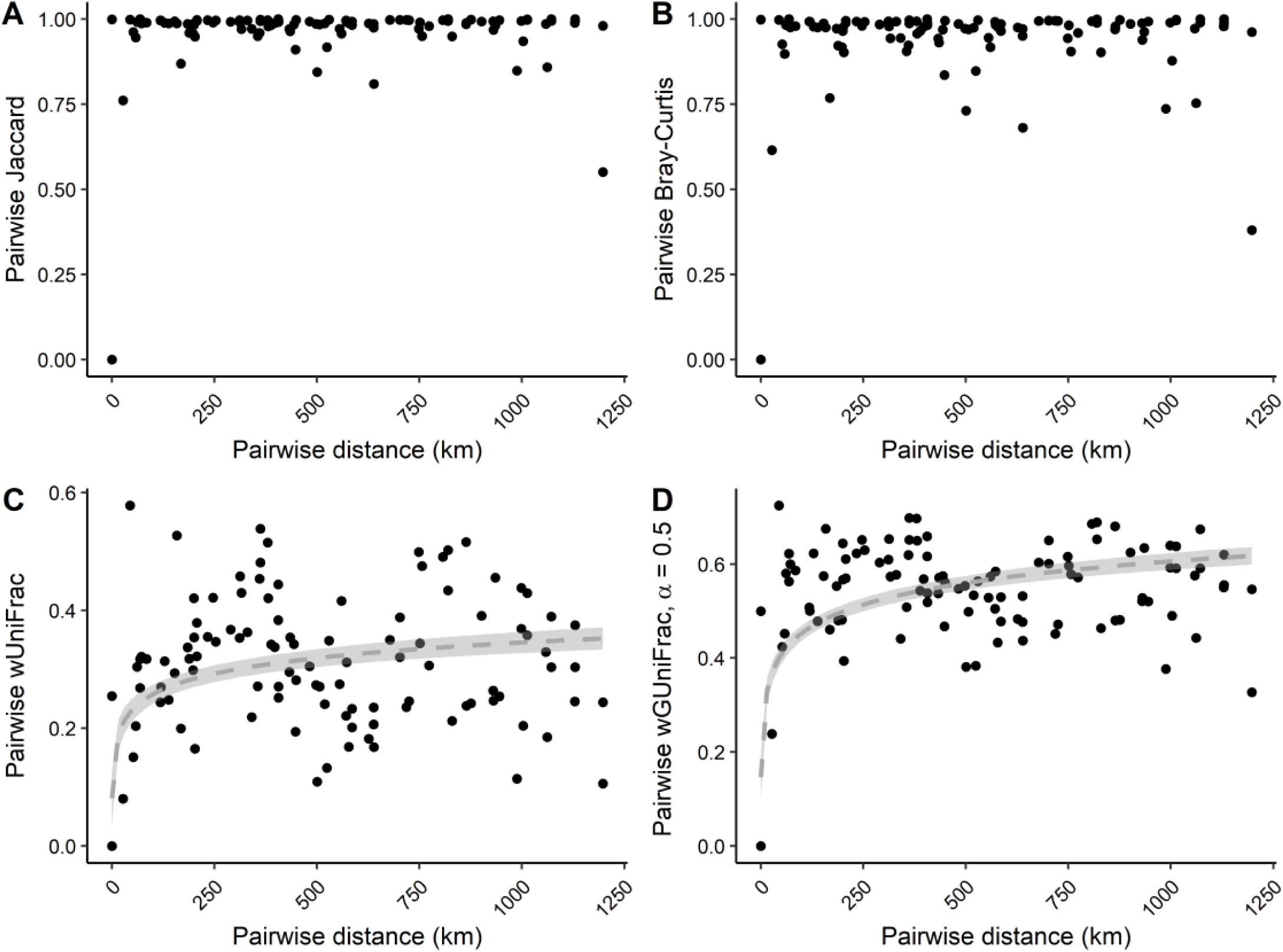
*Streptomyces* community pairwise dissimilarity metrics regressed onto pairwise geographic distance. The presence/absence formula for Jaccard distance (A) and abundance-weighted Bray-Curtis (B) metrics were calculated between pairs of sites and plotted against pairwise geographic distance (km). We also calculated dissimilarity metrics that incorporate phylogenetic information between samples using the original weighted UniFrac (wUnifrac) (C) and GUniFrac method (wGUniFrac) (D) that accounts for overweighting of proportional abundances of the original weighted Unifrac method (Wong *et al*., 2016). Grey dashed lines in panels C and D represent generalized additive models of the formula y ∼ ln(x + 1) and transparent grey shading surrounding the regression line represents upper and lower 95 % confidence intervals.

**Figure 3.**
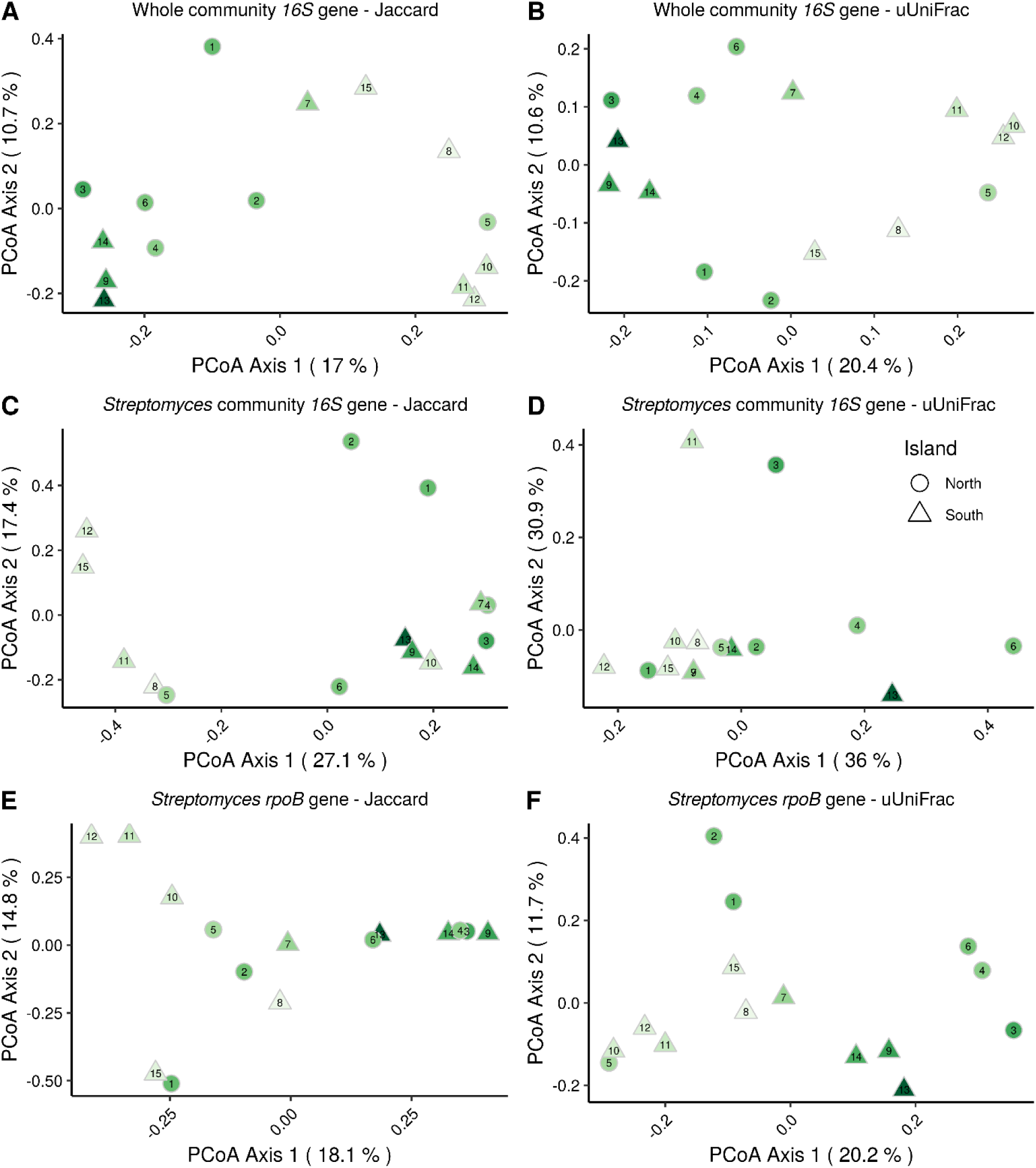
Multidimensional scaling (PCoA) ordinations of *Streptomyces rpoB* and 16S gene pairwise community dissimilarity among New Zealand sites. The proportion of variation explained by each coordinate axis is presented in parentheses on each axis. Sites located within New Zealand’s North or South Island are represented by circles and triangles, respectively. The green fill color represents pH, which ranges from 4.08 (light green) to 6.09 (dark green). Numbers inside symbols indicate the site number reported in Table 2 and Figure 4. The metric used to calculate pairwise dissimilarity is reported in the plot title. uUniFrac = unweighted UniFrac metric.

With respect to *Streptomyces* community composition, pairwise dissimilarity between sites saturated within 250 km (Fig. 2). Saturation of pairwise dissimilarity was less rapid for UniFrac metrics that incorporate phylogenetic distance (Fig. 2), suggesting phylogenetic conservation of community composition at regional spatial scales (< 250 km). The abrupt saturation in pairwise dissimilarity observed between sites is surprising considering *Streptomyces* form desiccation resistant aerial spores that facilitate dispersal (Sarmiento-Vizcaíno *et al*., 2016). However, dispersal depends on both displacement and colonization. The pattern of high dominance and high community dissimilarity that we observe suggests that priority effects and/or antagonistic interactions likely constrain colonization success despite high rates of spore displacement (Fukami, 2015; Wright and Vetsigian, 2016). Antagonism mediated by secondary metabolite production is common among divergent *Streptomyces* spp. (Traxler and Kolter, 2015), and such antagonistic interactions can promote coexistence of closely related species (Vetsigian *et al*., 2011; Cordero *et al*., 2012).

**Table 2.** Names and characteristics of soil sites sampled across New Zealand.

| Number | Code | Site | Latitude | Longitude | pH | MAR<br>(mm)<sup>a</sup> | MAT (°C)<sup>a</sup> | MAWS<br>(m/s)<sup>a</sup> | Elevation<br>(m)<sup>b</sup> |
| --- | --- | --- | --- | --- | --- | --- | --- | --- | --- |
| 1 | LB4N | Great Barrier Island 4N | -36.33 | 175.49 | 5.2 | 1294.8 | 14.3 | 4 | 541 |
| 2 | LB1B | Great Barrier Island 1B | -36.33 | 175.49 | 5.3 | 1294.8 | 14.3 | 4 | 541 |
| 3 | ACOR | Oteha Rowe | -36.73 | 174.69 | 5.5 | 1067.1 | 13.8 | 2.6 | 11 |
| 4 | NZOH | Ohinewai | -37.98 | 175.76 | 5 | 1018.1 | 14 | 1.8 | 130 |
| 5 | NZRD | Rangipo Desert | -39.4 | 175.8 | 4.8 | 2843.2 | 6.7 | 4.2 | 970 |
| 6 | NZMW | Mangaweka | -39.8 | 175.8 | 5.2 | 1134.7 | 11.1 | 2.3 | 357 |
| 7 | NZPA | Pangatotara | -41.18 | 172.9 | 5 | 1069.2 | 12.6 | 1.2 | 47 |
| 8 | NZHS | Hope Saddle<br>Lookout | -41.63 | 172.72 | 4.3 | 2906.2<sup>c</sup> | 12.2<sup>c</sup> | 3.5<sup>c</sup> | 603 |
| 9 | NZKP | Kaikoura Peninsula | -42.42 | 173.7 | 5.6 | 1031.2<sup>c</sup> | 12.2 | 4.3 | 70 |
| 10 | NZWS | Woodstock Road | -42.82 | 170.9 | 4.5 | 2197.2 | 12.2 | 3.8 | 22 |
| 11 | NZHV | Haast Valley | -43.94 | 169.15 | 4.6 | 2906.2 | 11.3 | 2.8 | 29 |
| 12 | NZCF | Cameron Flats | -44.16 | 169.29 | 4.4 | 727.6 | 9.2 | 2.7 | 346 |
| 13 | NZKG | Kawarau Gorge | -44.7 | 169.13 | 6.1 | 454.6 | 10.6 | 2.8 | 287 |
| 14 | NZKR | Kingsland Rd | -45.15 | 168.75 | 5.5 | 735.3 | 9.7 | 3.5 | 310 |
| 15 | NZRT | Red Tussock<br>Conservation Area | -45.56 | 168.04 | 4.4 | 1031.2 | 9.5 | 2.6 | 486 |

<sup>a</sup>Mean annual rainfall (MAR), temperature (MAT), and wind speed (MAWS) data were accessed from the National Institute of Water and Atmospheric Research (NIWA) CliFlo (https://cliflo.niwa.co.nz/) database weather station locations nearest ( $35 \pm 21$ km) to sites where soils were sampled.
<sup>b</sup>Elevation was accessed from https://www.maps.ie/coordinates.html.
<sup>c</sup>Values were imputed using the R package mice (see Materials and Methods for details).

In addition to signatures of dispersal limitation, we detected a positive association between *Streptomyces* community net relatedness index (NRI) and latitude (F_1,13_ = 3.59; Adj. R^2^ = 0.16; *p* = 0.08; Fig. 4A-B), with NRI values increasing towards the equator. The NRI is a standardized measure of phylogenetic clustering and increases or decreases when phylogenetic diversity is lower (clustered) or higher (over-dispersed) than expected considering the species richness (Webb *et al*., 2002; Swenson, 2009). Although this relationship was not significant, removal of three suspected outliers (points 2, 14, 15; Fig. 4A) revealed a significant relationship (F_1,10_ = 9.03, *P* = 0.013, Adj. R^2^ = 0.42) between NRI and latitude. A significant NRI-latitude relationship was observed for *Streptomyces* communities in North America, with phylogenetic clustering increasing poleward (Andam *et al*., 2016). However, the NZ NRI-latitude association is opposite to the previously described trend, with phylogenetic clustering increasing towards the equator (Fig. 4A-B).

**Figure 4.**
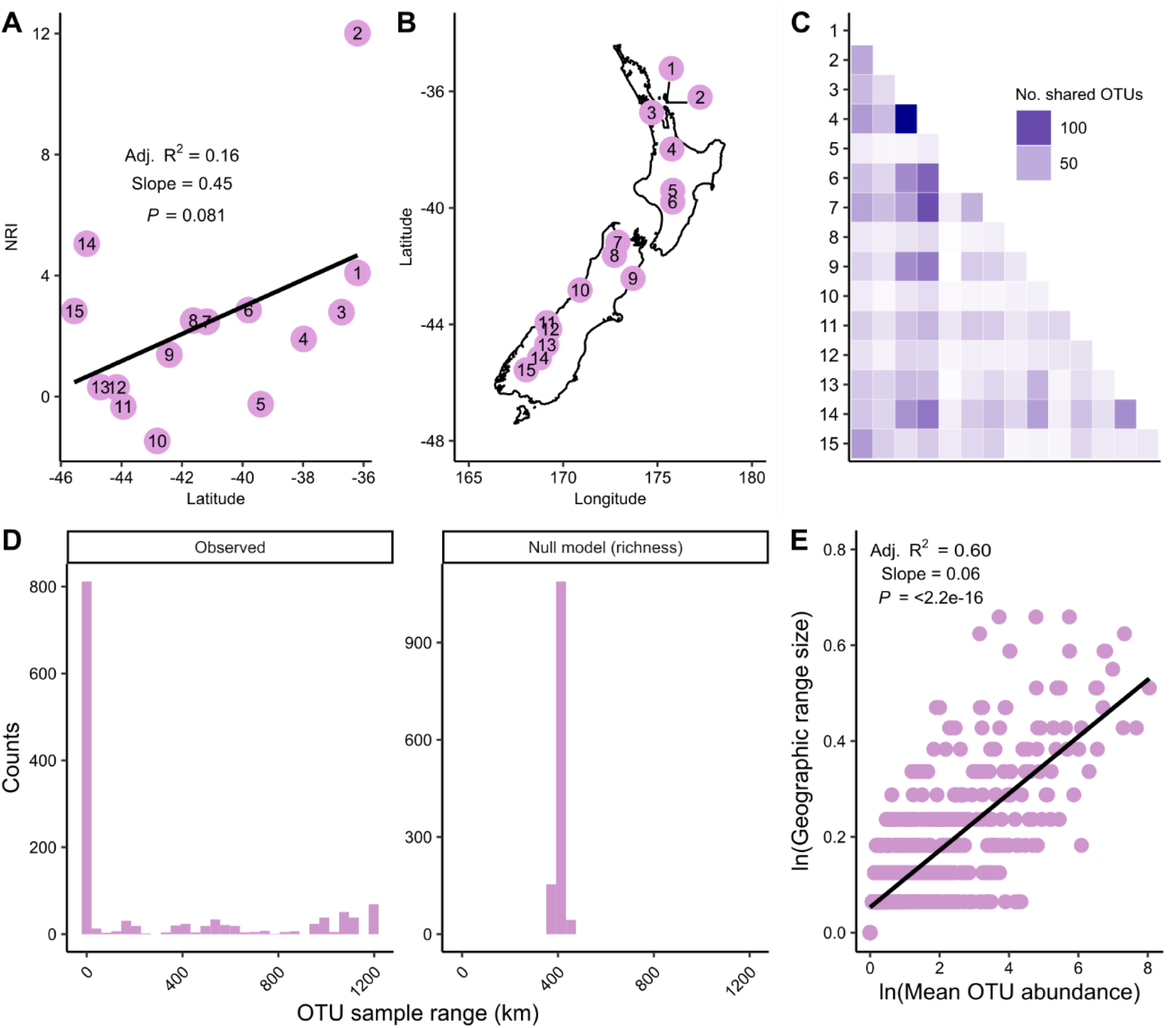
Distribution and diversity of *Streptomyces rpoB* OTUs in soils along a latitudinal transect across New Zealand. Net relatedness index (NRI) regressed on latitude (A) across the 15 study sites in New Zealand (B). The pairwise number of shared *rpoB* OTUs among sites (C). The distribution of range (km) values for each *rpoB* OTU for the observed and null model data sets (D). The relationship between natural log transformed mean *Streptomyces* OTU relative abundance and proportion of occupied sites (E). OTU sample range in the richness null model panel represents the average of 1,000 null model iterations.

The *Streptomyces* NRI-latitude relationship described by Andam et al. is consistent with niche conservatism following retreat of continental glaciers in North America after the last glacial maximum (LGM) (Andam *et al*., 2016). However, the LGM in NZ (∼25-10 kya) was characterized by montane glaciers that were largely restricted to discrete regions of NZ’s south island (McGlone, 1985; Wallis and Trewick, 2009; Newnham *et al*., 2013). Glacial dynamics in New Zealand during the LGM differed from those of continental regions. Recolonization dynamics in NZ at the end of the LGM are hypothesized to have resulted from species expansion from scattered ‘micro-refugia’ rather than northward expansion from southern refugia as hypothesized for North America and Europe (Provan and Bennett, 2008; Wallis and Trewick, 2009). Furthermore, the predicted vegetation cover of ice-free regions on NZ’s South Island during the LGM was predominantly dry grassland and shrubland habitat in which *Streptomyces* species thrive (Newnham *et al*., 2013; Kämpfer *et al*., 2014), whereas forested habitat, within which *Streptomyces* are not favored, was more common on the North Island (McGlone, 1985; Newnham *et al*., 2013). The climatic stability of habitat favorable to *Streptomyces* on the South Island (De Wever *et al*., 2009; Keppel *et al*., 2012) may explain the high phylogenetic diversity (lower NRI values) of sites in southern NZ (Fig. 4A). In contrast, the persistence of beech and podocarp forests on the North Island may have selected specifically for *Streptomyces* species capable of degrading plant woody components (Bontemps *et al*., 2013), resulting in phylogenetically clustered communities (positive NRI values, Fig. 4A). These results are consistent with the hypothesis that historical changes in paleoclimate have influenced extant patterns of *Streptomyces* biogeography.

Patterns of *rpoB* OTU sharing among sites revealed greater *rpoB* OTU sharing between North and South Island sites than within comparisons of South Island sites alone (Fig. 4C), which is consistent with a legacy of glaciation on NZ’s South Island. Some South Island sites exhibited low *rpoB* OTU sharing (sites 5, 8, 10, 12; Fig. 4C), and three of these four sites have roughly double the estimated MAR of other sites (Table 2), suggesting environmental filtering could be contributing to the observed co-occurrence patterns. We also investigated *rpoB* OTU ranges within NZ, which were observed to be highly constrained for most OTUs (Fig. 4D, observed panel). To assess whether the observed species ranges were an artifact of the sampling design, we randomly shuffled *rpoB* community data using a null model algorithm that constrains site richness to the observed richness but enables *rpoB* OTU occupancy to vary freely between sites (Fig. 4D, richness panel). Under this null model scenario, the average *rpoB* OTU range (408 km) approaches the average pairwise distance among sites (496 km), which deviates considerably from the observed *rpoB* OTU range distribution and indicates species distributions are not random. While a significant fraction of *Streptomyces* OTUs possess a wide distribution (13% of *Streptomyces* OTUs have sample ranges > 1,000 km; Fig. 4D), the majority appear to be constrained by biotic or abiotic filters that impair their recruitment to only one or a few local sites across NZ. Furthermore, we observed a positive relationship between *Streptomyces* OTU mean relative abundance and their geographic range size (Fig. 4E). This relationship has been documented for bacteria, plants, and animals (Brown, 1984; Gaston *et al*., 1997; Harte *et al*., 2001; Nemergut *et al*., 2011), but information regarding range size of discrete bacterial lineages using high-resolution genetic markers are lacking. Multiple mechanisms to explain this relationship have been hypothesized, such as neutral processes, differential dispersal capabilities (e.g., spore morphology differences among *Streptomyces* OTUs), differential birth and death rates, habitat selection, and differential resource utilization (Gaston *et al*., 1997; Nemergut *et al*., 2011), but further investigation is required to identify the underlying causes of the abundance-range relationship for *Streptomyces* OTUs.

The variables pH, MAR, and plant species richness were all significantly associated with *Streptomyces* community composition (jaccard-based PCoA; envfit R^2^ = 0.49, 0.44, 0.64 and *p* = 0.016, 0.029, and 0.002, respectively; Table S2). Similar results were observed using an unweighted UniFrac PCoA ordination (Table S2). Accordingly, pH was significantly associated with Jaccard-based PCoA axis 1, whereas PCoA axis 2 was significantly associated with NZ plant species richness (Fig 5A, 5B, respectively). Although the association between NZ plant species richness and the Jaccard-based ordination remained significant when controlling for both MAT, MAR, and latitude (t_3,11_ = −2.71, *p* = 0.022), our sample size is limited and our results should be viewed as preliminary. Regardless, this finding warrants further study due to the plant-associated lifestyle of many members of the *Streptomyces* and prior reports of links between *Streptomyces* community composition and plant diversity (Bakker *et al*., 2013; Viaene *et al*., 2016; Rey and Dumas, 2017).

**Figure 5.**
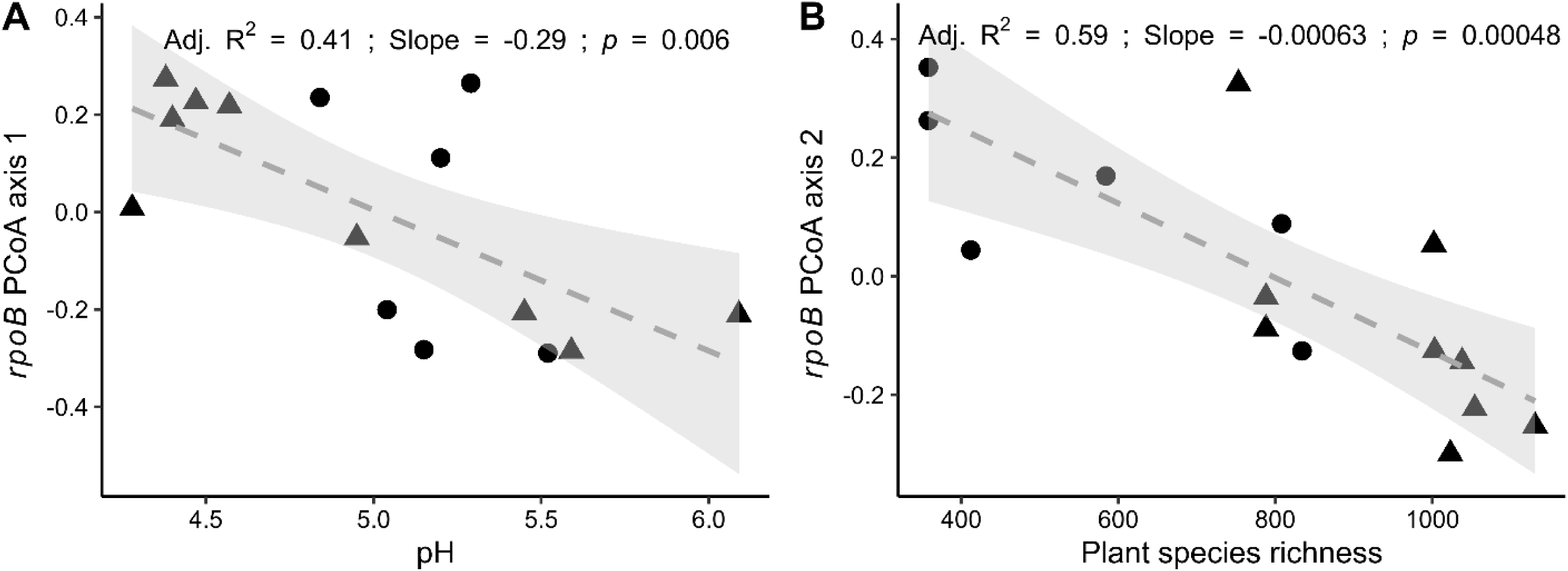
The association of *Streptomyces* community beta diversity with pH and NZ plant species richness within 100 km of the NZ sites. The first and second ordination axes of a Jaccard-based *Streptomyces* community PCoA were regressed against pH (A) and plant species richness (B).

#### Community assembly processes governing Streptomyces community turnover

Variable selection was identified as the greatest driver of community assembly across the NZ sites (Table 3), while the contribution of homogeneous selection to community turnover was low (7% of all comparisons). These results are not surprising considering the geographic scale of our analysis, which encompasses sites with widely varying environmental characteristics (Barker, 2005). Homogenizing dispersal contributed to approximately 30% of community assembly processes identified (Table 3). Most pairwise site comparisons within 500 km had β_RC_ values < – 0.95, which indicates greater community similarity than expected when compared with null modelling (Fig. S4A). However, some β_RC_ values < −0.95 were also detected between sites located >1,000 km apart, which may be indicative of long range dispersal of *Streptomyces* spp. (Fig. S4A). The potential for long-range dispersal of *Streptomyces* has been observed previously (Andam *et al*., 2016; Choudoir *et al*., 2016), and 13% (n = 171) of the *rpoB* OTUs were identified in New Zealand sites separated by >1,000 km. While dispersal limitation accounted for only 6% of community assembly patterns, values of β_RC_ increased with increasing pairwise distance and suggests an underestimation of dispersal limitation in our analysis (Fig. S4B). One possible explanation is that non abundance weighted metrics may not achieve the same sensitivity as their abundance weighted counterparts (Stegen *et al*., 2012, 2015). However, abundance weighting of β_RC_ as RC_bray_ (Stegen *et al*., 2013) is inappropriate for dry-stored soils for which sample abundance might not represent *in situ* abundance (Clark and Hirsch, 2008; Ivanova *et al*., 2017). While the β_RC_ metric is also influenced by the size of the regional species pool selected (Chase *et al*., 2011), *Streptomyces* spores can be readily dispersed by insects or wind (Lloyd, 1969; Ruddick and Williams, 1972), and the choice of the totality of NZ as a regional *Streptomyces* species pool is not unrealistic. Regardless, the approach that we have applied, combining null modeling with multiple lines of evidence, has been shown to provide robust insight into assembly processes governing microbial communities (Chase *et al*., 2011; Stegen *et al*., 2015).

**Table 3.** List of community assembly processes and their overall contributions to *Streptomyces* community dynamics.

| Process | No. pairwise comparisons | Percentage of total |
| --- | --- | --- |
| Variable selection | 72 | 34 |
| Homogeneous selection | 14 | 7 |
| Dispersal limitation | 12 | 6 |
| Homogenizing dispersal | 62 | 29 |
| Undominated<sup>a</sup> | 50 | 24 |

<sup>a</sup>Neither selection nor dispersal dominates.

### Comparison of environmental rpoB to cultivated representatives

#### Streptomyces rpoB OTUs and uncultivated rpoB sequence diversity

Of the *rpoB* OTUs classified as *Streptomyces*, only 12% (n = 149) matched *rpoB* sequences from cultivated *Streptomyces* strains (n = 1808) described in the ARS Microbial Genomic Sequence Database (99% nucleotide identity cutoff; see Materials and Methods for details). Therefore, despite their cultivability (Kämpfer *et al*., 2014), only a small fraction of *Streptomyces rpoB* OTUs were linked to cultivated representatives. However, the 149 OTUs linked to cultivated strains represented one quarter of all sequence counts (108,068/428,122), and four of these OTUs (OTU_3, OTU_5, OTU_6, and OTU_8) represented 65% (n = 70,652) of this fraction. In addition, the percentage of *rpoB* OTUs matching *rpoB* sequences extracted from 998 *Streptomyces* genomes was only 10% (n = 126) of the 1,287 OTUs observed. Of the four dominant cultivated OTUs, OTU_6 failed to match an *rpoB* sequence from the 998 genomes examined. The *rpoB* sequence of OTU_6 was most similar to ARS type specimen *Streptomyces sanglieri* strain NRRL B-24279 (99.7% nucleotide identity), but was only 98.857% similar to the genome of *Streptomyces* sp. DvalAA-43 (NCBI accession FMBW01000037.1), suggesting a representative genome for this strain may be lacking. Thus, data derived from the *rpoB* primer set may assist in prioritizing future culture collection sequencing efforts of *Streptomyces* isolates. The fact that *rpoB* OTUs matching cultivated strains represented 12% of taxa observed but 25% of sequence counts suggests that most *Streptomyces* communities are dominated by few taxa and contain many rare taxa. Since isolation of many pure cultures is laborious, isolation efforts are likely to under sample *Streptomyces* diversity. In addition, the low representation of *rpoB* OTUs among cultivated strains and sequenced genomes suggests that the genus *Streptomyces* contains an enormous genomic diversity that remains largely uncharacterized. Future cultivation efforts might combine high-throughput *Streptomyces rpoB* amplicon sequencing and single cell (or spore) cultivation to capture rare taxa present in *Streptomyces* communities.

## Conclusions

Biogeographical surveys of the *Streptomyces* have long benefitted from cultivation-dependent methodologies, but the development of efficient, cultivation-independent methodologies are preferred for large scale biogeographic studies. We introduce a high-throughput sequencing primer set that targets the DNA-directed RNA polymerase beta subunit (*rpoB*) within the bacterial genus *Streptomyces* and examined the biogeography of *Streptomyces* in soils across a 1,200 km transect in New Zealand (NZ). The *rpoB* PCR primers provided a near complete inventory of *Streptomyces* OTUs in NZ soils and provided far greater coverage of *Streptomyces* spp. than could be achieved by cultivation efforts or by 16S rRNA gene sequencing data from the same sites. Specific abiotic (pH, MAR), biotic (plant richness), and historical (glaciation) features of NZ were found to significantly correlate with *Streptomyces* PD. Phylogenetic clustering of *Streptomyces* communities was opposite to that observed in North America, with clustering increasing towards the equator. Presumably, the presence of favorable grassland and shrubland refugia afforded by restricted, montane glaciers in NZ contributed to the lack of phylogenetic clustering on NZ’s South Island. These results, taken together, suggest that extant *Streptomyces* communities are shaped by dispersal of spores from species pools determined by historical biogeographical processes, with community structure of individual sites shaped by competitive interactions that govern colonization dynamics and which are constrained by local abiotic (soil pH, temperature) and biotic (plant communities) conditions. Overall, the *rpoB* primer set enabled the detection of a heretofore unobserved *Streptomyces* diversity, greatly exceeding those currently documented in culture and genome collections, and hinting at an untapped reservoir of novel *Streptomyces* antibiotics and biologics.

## Experimental Procedures

### Soil sampling, storage, and data acquisition

Soils (0-10 cm depth) were sampled from 15 locations up to ∼1,200 km apart throughout NZ between February and April of 2012 (Table 2). Soils were sieved (2 mm mesh), air-dried, and stored in sterile, plastic whirl-pak bags (Nasco, Fort Atkinson, WI, USA) at room temperature until DNA extraction (∼ 3 yr). Air-dried, archived soils have been extensively studied and represent a valuable resource for soil microbiologists due to the cost and space required to maintain frozen soil samples (Cary and Fierer, 2014; Dolfing and Feng, 2015). However, the community composition of archived, air-dried soils is significantly different from frozen/fresh soils and careful consideration is required when analyzing air-dried soil that has been stored for long periods (years) (Dolfing *et al*., 2004; Clark and Hirsch, 2008; Tzeneva *et al*., 2009; Ivanova *et al*., 2017). *Streptomyces* produce desiccation resistant spores that can survive dry conditions for many years (Ensign, 1978). While we found that storage method did alter *Streptomyces* relative abundance, analysis of air dried relative to frozen soils indicated significant correlation in the relative abundance of *Streptomyces* OTUs between the two storage methods (see SI and Fig. S3 for additional details).

The pH of the soil was measured by pH probe using 1:2 (w/v) soil dilutions in 10 mM CaCl_2_ (Staff, 2014). The mean annual air temperature (MAT) (°C), rainfall (MAR) (mm), and wind speed (MAWS) (m/sec) for 2012 were retrieved from the National Institute of Water and Atmospheric Research (NIWA) CliFlo (https://cliflo.niwa.co.nz/) database weather stations nearest (35 ± 21 km) to soil sampling sites. Plant species survey data (NZ National Vegetation Survey databank; https://landcareresearch.co.nz/) were accessed from the Global Biodiversity Information Facility (https://www.gbif.org/) in December 2018. Only species presence/absence data from vegetation plots surveyed after the year 1960 and within 100 km of the soil sampling sites were used as input for downstream analyses. To control for sampling effort, plant species presence/absence data were generated by rarefying to the minimum number of observations (9,790) per site and resampling with replacement 999 times. The overall presence/absence of each plant species was scored at a site by iterating over all species identified in the dataset and scoring their presence/absence within all 999 draws of 9,790 observations for each site (a link to a figshare repository containing the raw data is provided in the *Data availability* section).

### Microbial strains for PCR amplification

Genomic DNA stocks prepared from *Streptomyces* isolates cultivated from a variety of habitats were used as positive controls for *rpoB* amplification (Choudoir *et al*., 2016). Additional isolates of *Streptomyces, Kitasatospora, Streptacidiphilus*, and *Catenulispora* were received by request from the United States Department of Agriculture Agricultural Research Service culture collection (https://nrrl.ncaur.usda.gov). These species were cultivated as indicated by documentation accompanying the isolate and DNA was extracted by one minute of bead-beating (0.1 mm silica beads) using an established chloroform-phenol extraction method (Griffiths *et al*., 2000).

### PCR primer design and analysis, DNA isolation, and Illumina sequencing

*Streptomyces rpoB* gene sequences were downloaded from the PubMLST database (Jolley *et al*., 2018), nucleotide alignment performed with MAFFT, and PCR primers designed using DECIPHER software (Wright, 2016) to amplify a region of sufficient length for successful Illumina high-throughput amplicon sequencing projects (usually ∼200-500 nt). DECIPHER was also provided *rpoB* sequences from members of the *Kitasatospora* (N = 10) and *Streptacidiphilus* (N = 9) to enable DECIPHER to discriminate against these sequences during primer design and enhance specificity of the primers for *Streptomyces rpoB* gene sequences (see SI Results and Discussion for additional details). The *rpoB* primers were named Smyces _rpoB1563F (5’-GGAGGACCGCTTCGTCATC-3’) and Smyces_rpoB1968R (5’-GTACGTGGTGTACGTGCCG-3’) and amplify a 406 bp region within the *rpoB* gene. The numbers within the primer names correspond to the primer alignment position within the *rpoB* sequence from *Streptomyces coelicolor* strain A3(2) (NCBI accession NP_628815.1). The *rpoB* primers containing modifications for Illumina sequencing (as per (Kozich *et al*., 2013)) are provided in Supporting Information (SI) (Table S3).

Forward and reverse primers targeting the V4 region of the 16S rRNA (16S) gene in bacteria (5’-GTGCCAGCMGCCGCGGTAA-3’ and 5’-GGACTACHVGGGTWTCTAAT-3’, respectively) were used for overall bacterial community analyses in NZ soils as previously described (Kozich *et al*., 2013). DNA was extracted from NZ soils using a PowerSoil DNA isolation kit (Qiagen, Hilden, Germany) following manufacturer’s protocols and subsequently stored at −20 °C until use. Briefly, *rpoB* and 16S gene end-point PCR reactions consisted of 1 µl of DNA (∼0.5-19.5 ng DNA total), 12.5 µl of Q5 Hot Start High-Fidelity 2X Master Mix (New England Biolabs, Ipswich, MA, USA), 0.625 µl of 4X Quant-iT PicoGreen dsDNA assay reagent (Thermo Fisher Scientific, Waltham, MA, USA), 1.25 µl each of 10 µM dual-barcode forward and reverse primers, 1.25 µl of 20,000 ng/µl bovine serum albumin (New England Biolabs), and 7.125 µl of molecular grade water. Thermocycler conditions were 98 °C for 30 sec, 30 cycles of 98 °C for 10 sec, 55 °C for 30 sec, and 72 °C for 1 min, followed by extension for 5 min and melt curve analysis at 0.5 °C/5 sec from 65 to 95 °C. PCR Amplification of *rpoB* gene amplicons from a variety of *Streptomyces* isolates revealed clean, consistent amplicons of the expected size (Fig. S5). Although *Streptomyces rpoB* gene copies per g soil were not quantified in the present study, the *rpoB* primer pair reported can also be used in quantitative PCR assays. The details of these methods are reported in SI (Table S4, Fig. S6).

After PCR amplification, all end-point *rpoB* and 16S gene amplicons were pooled at equimolar concentrations (∼1-2 ng/µl) using a SequalPrep normalization kit (Thermo Fisher Scientific, Waltham, MA, USA) and purified by gel extraction using the Wizard SV gel and PCR clean-up system (Promega, Fitchburg, WI, USA) following manufacturer’s instructions. The resulting sequencing libraries were provided to the Cornell University Biotechnology Resource Center for paired-end Illumina MiSeq sequencing (2 × 250 bp) (Illumina, Inc., San Diego, CA). A 20 pM concentration of phiX spike-in at 10-20 % [v/v] was added to 16S and *rpoB* gene sequencing libraries to improve the quality of the low diversity amplicon libraries.

### Sequence processing and analysis

For simplicity of reporting our 16S gene processing, we report both OTU and exact sequence variant (ESV) identification methods for the *rpoB* gene. However, we only present OTU data for the *rpoB* gene in the manuscript. While ESVs are recommended for analyses of microbial communities using the 16S gene (Knight *et al*., 2018), their utility for protein-coding nucleotide sequences is uncertain considering distinct ESVs could comprise synonymous or non-synonymous substitutions occurring within a single species. Furthermore, empirical analysis demonstrated that 99% nucleotide identity OTUs are a reasonable proxy for species when compared with a species definition for *Streptomyces* using multi-locus sequence analysis (Fig. S7) (Rong and Huang, 2012). See SI for details regarding the analysis.

Raw sequences were demultiplexed using dual-barcode FASTQ sequence information (Kozich *et al*., 2013) with the tool deML (Renaud *et al*., 2015), followed by paired-end read merging using PEAR v0.9.10 (with parameters -p 0.001 and –q 30) (Zhang *et al*., 2014). Residual Illumina adapters and *rpoB* primer sequences were removed with Cutadapt v1.14 (Martin, 2011). The merged *rpoB* reads were either clustered *de novo* at 99% nucleotide identity or identified as exact sequence variants (ESVs) with VSEARCH v2.8.1 (Rognes *et al*., 2016) or USEARCH (UNOISE3 function) v10.0.240 (Edgar, 2016), respectively. Prior to 99% *rpoB* OTU clustering with VSEARCH, *rpoB* sequences were filtered using a per base expected error of 0.25 (--fastq_maxee 0.25) and trimmed to 350 nt in length (--fastq_trunclen 350). The output sequences were dereplicated (singletons removed), pre-clustered at 99.5% nucleotide identity, and chimeric sequences identified and removed by UCHIME (--uchime_denovo option in VSEARCH) (Edgar *et al*., 2011). For ESV identification, UNOISE3 was run using default settings except the minimum abundance was set to 9 (-minsize 9) to filter out ESVs that may represent sequencing artifacts. Taxonomic assignments were made using the BLCA tool (Gao *et al*., 2017) (using parameters -n 100 -j 50 -d 0.1 -e 0.01 -b 100 -c 0.9 --iset 90) and a custom *rpoB* sequence database (see below). The classification of 16S gene OTUs and ESVs followed a similar methodology, except that a 97% nucleotide identity cutoff was provided to VSEARCH for 16S gene OTU identification and taxonomy assigned using BLCA and NCBI’s 16S rRNA gene collection (Gao *et al*., 2017). The number of *rpoB* reads per sample varied between 925 and 90,276 (25,658 ± 26, 732, mean ± s.d.). The number of reads per sample for 16S gene OTUs and ESVs ranged between 12,574 and 45,532 (21,529 ± 8,803, mean ± s.d.) and 16,974 and 55,920 (27,328 ± 10,640), respectively. Of the 3,322 *rpoB* OTUs detected in the present study, only 1,297 (39%) were classified as members of the *Streptomycetaceae* (Table 4). However, the abundance of the 2,025 non-target OTUs comprised only 9% (39,866 / 467,988) of the total read counts (Table 4; see SI for additional details).

**Table 4.** Proportion of *rpoB* OTUs classified as *Streptomyces* spp. or other taxa.

| Taxon | No. OTUs | Percentage of reads |
| --- | --- | --- |
| *Streptomyces* | 1,287 | ~91% |
| Non-*Streptomyces* within *Streptomycetaceae* | 10 | <1% |
| Non-target<sup>a</sup> | 2,025 | ~9% |

<sup>a</sup>Includes *rpoB* and *rpoC* gene sequences.

A phylogeny of 16S gene and *rpoB* OTU sequences was generated by sequence alignment with MAFFT v7.407 (Katoh *et al*., 2009) followed by tree construction using FastTree v 2.1.8 (Price MN, Dehal PS, 2010) (with parameters -gtr -gamma -spr 4 -mlacc 2 -slownni – bionj –nt). The resultant phylogeny was midpoint-rooted with the R package phangorn (Schliep, 2010). Visual inspection of the *rpoB* gene phylogeny revealed three OTUs classified as *Streptomyces* (OTU_3178, OTU_3273, and OTU_3090) possessing long branch lengths of 6.0611, 0.1380, and 6.0579, respectively, and low sequence counts (≤ 5 counts). As a reference, the interquartile branch length range was between 0.0033 and 0.0134. Hence, the three *rpoB* OTUs were removed prior to analysis since they may represent sequencing artifacts.

Available *Streptomyces* genomes were accessed from NCBI (as of 9/11/2017; n = 998) and prodigal v2.6.2 (Hyatt *et al*., 2010) was used to predict protein-coding gene and amino acid sequences for consistent gene prediction among genomes. Then, predicted protein-coding sequences were provided to CheckM v1.0.7 (Parks *et al*., 2015) via the analyze function and the output was parsed for sequences containing an N-terminal protein domain diagnostic of bacterial RpoB (RNA_pol_Rpb2_1, pfam id: PF40563). In total, 990 full-length *rpoB* sequences were used as a reference genome database. An additional set of 1,834 partial *rpoB* gene sequences from cultivated *Streptomyces* species were accessed from the ARS Microbial Genomic Sequence Database server (http://199.133.98.43) (Labeda *et al*., 2017). The nucleotide version of the basic local alignment search tool (BLAST) (Altschul *et al*., 1990) was used to align *rpoB* OTUs to reference *rpoB* sequences from both genomes and type culture collections. An *rpoB* OTU was considered as a ‘match’ to a reference *rpoB* sequence when alignments satisfied the criteria of ≥ 99% nucleotide identity and ≥ 99% query coverage. Of the 1,834 partial *rpoB* database sequences, 85 were identified as *Streptomyces* species associated with scab-causing disease (Zhang *et al*., 2016). Any *rpoB* OTUs demonstrating significant alignment (≥ 99% nucleotide identity, ≥ 99% query coverage) to these sequences were classified as putative pathogens.

### Statistics

The R package phyloseq was used to pre-process or manipulate 16S and *rpoB* community data (McMurdie and Holmes, 2013). The iNEXT package was used for calculating richness estimates and rarefaction of the *rpoB* community data (Hsieh *et al*., 2016). The *rpoB* OTU richness estimates were normalized to a consistent sample coverage among samples (95% in the present study; Chao et al. 2014; Mateo-Tomás et al. 2015) when comparing *rpoB* OTU richness to spatial (latitude), environmental (pH, temperature, rainfall, wind speed), and biological (plant species diversity) correlates of *Streptomyces rpoB* OTU richness in NZ. Indirect gradient analysis of the *rpoB* dissimilarity data was performed using multidimensional scaling (PCoA) with the cmdscale command in R (R Core Team, 2018). The adonis and envfit functions (Oksanen *et al*., 2019) were used to evaluate whole community associations with environmental variables using 9,999 permutations. Ordinary least squares regression was performed with the R ‘stats’ package and regression tables produced using the R package ‘stargazer’ (Hlavac, 2015; R Core Team, 2018). Community evenness was estimated using the codyn package in R and the E_var_ evenness metric, which ranges between 0 (complete dominance) and 1 (complete evenness) (Smith *et al*., 1996; Hallett *et al*., 2016). We estimated species contributions to beta diversity (SCBD) from the Hellinger-transformed *Streptomyces* OTU table using the beta.div function in the adespatial R package (Dray *et al*., 2017). The net relatedness index (NRI), an estimate of phylogenetic diversity compared to a null model of species richness, was estimated using the picante package and multiplied by −1 for comparability with previous phylocom NRI values using *Streptomyces* community data (Kembel *et al*., 2010; Andam *et al*., 2016). Missing NIWA climate values (rainfall and temperature for Hope Saddle Lookout and rainfall for Kaikoura Peninsula) were imputed with the R package ‘mice’ using the ‘pmm’, or predictive mean matching, method (van Buuren and Groothuis-Oudshoorn, 2011). We investigated the contributions of selection, dispersal, drift using a null modeling approach (*sensu* Stegen) by comparing non-abundance weighted (presence-absence) beta mean nearest taxon index (βNTI) and the beta Raup-Crick metrics (β_RC_) (Chase *et al*., 2011; Stegen *et al*., 2013, 2015). The original β_RC_ metric was preferred over the RC_bray_ metric since the latter requires relative abundance information that may not be appropriate for air-dried, archived soils. Null modeling of βNTI was performed by subtracting from the observed beta mean nearest taxon distance (βMNTD) the mean of the null βMNTD and dividing by the s.d. of the null βMNTD (999 replicates, taxon labels shuffled using tipShuffle in the R picante package) (Kembel *et al*., 2010; Stegen *et al*., 2013; Swenson, 2014). Values of β_RC_ were calculated using 9,999 replicates with the raup_crick function reported by Chase et al. (2011). The simultaneous analysis of both βNTI and β_RC_ metrics enables the partitioning of community variation into components of selection (variable and homogeneous, βNTI > 2 and < −2, respectively), dispersal (limited and homogenizing dispersal, |βNTI| < 2 and β_RC_ > 0.95 or |βNTI| < 2 and β_RC_ < −0.95, respectively) and drift (neither selection nor dispersal dominated or “undominated”, |βNTI| < 2 and |β_RC_| < 0.95) (Stegen *et al*., 2015). Additional null modeling was performed with 1,000 iterations using the randomizeMatrix function in the picante package in R (Kembel *et al*., 2010).

## Data availability

Both 16S and *rpoB* gene sequences amplified from the NZ soil DNA were submitted to the Sequence Read Archive (Leinonen *et al*., 2010) under study no. SRP194926. The National Center for Biotechnology Information BioProject and BioSample accession numbers are PRJNA540985 and SAMN11569796, respectively. A figshare repository containing all scripts, data, and reference *rpoB* sequences is publicly accessible at (https://doi.org/10.6084/m9.figshare.c.4573250).

## Acknowledgements

S.A.H. is grateful to Dr. Ashely M. Frank for her assistance with discussions regarding manuscript writing, organization, and revision. D.H.B. is grateful to Dr. Paul B. Rainey and The Institute for Advanced Study, Massey University for providing research support, laboratory access, and helpful advice. This material is based upon work supported by the National Science Foundation under Grant No. DEB-1456821 awarded to D.H.B.

## Author Contributions

D.H.B collected samples and designed the experiment, S.A.H. and K.P. performed analyses, and S.A.H. and D.H.B. wrote the manuscript.

