## Supporting information for "The biogeography of *Streptomyces* in New Zealand enabled by high-throughput sequencing of genus-specific *rpoB* amplicons"

### Materials and Methods

*Effects of frozen vs dry soil storage on Streptomyces communities*

The community composition of archived, air-dried soils has been found to differ significantly from frozen/fresh soils and careful consideration is required when analyzing air-dried soil that has been stored for long periods (years) (Dolfing *et al.*, 2004; Clark and Hirsch, 2008; Tzeneva *et al.*, 2009; Ivanova *et al.*, 2017). Members of the *Streptomyces* possess a complex developmental cascade that terminates in the production of durable spores which maintain their viability when exposed to conditions that vegetative hyphae cannot survive (McBride and Ensign, 1987; Claessen *et al.*, 2006). To assess the effect of long-term storage (years) of air-dried versus frozen soil samples on the composition and diversity of *Streptomyces* species present in soil, we compared *Streptomyces rpoB* OTU abundances in forest and plantation soil stored frozen or air-dried for ~3 years. Although there were clearly departures from the 1:1 ratio of air-dried to frozen *rpoB* OTU abundances (Fig. S3), we observed a significant spearman rank correlation in *Streptomyces rpoB* OTU relative abundances in frozen and air-dried samples from both forest ($r_s = 0.45$, $P < 2.2e-16$) and plantation ($r_s$ =0 .12, $P = 0.001$) soils. Of note, log- and Hellinger-transformed OTU abundances marginally improved the rank correlation in OTU abundance between air-dried and frozen plantation soil ($r_s$ =0 .17, $P = 1.2e-5$), but not in forest soil ($r_s = 0.44$, $P < 2.2e-16$), suggesting that some of the effect of air-drying can be controlled by data transformation. Nonetheless, we largely focus on presence/absence alpha and beta diversity characteristics between NZ sites in the present study to minimize type I errors induced by soil storage (i.e., false positive associations). Since the rate of DNA decay is low in dry soils and a DNA signature in soil can last for years (Levy-Booth *et al.*, 2007; Sirois and Buckley, 2019), sufficient evidence exists to justify the analysis of presence/absence OTU data for long-term soil storage archives.

*Quantitative PCR*

The *rpoB* quantitative PCR reactions consisted of 2 µl genomic DNA, 10 µl of 2X SsoAdvanced Universal SYBR Green Supermix (Bio-Rad Laboratories, Hercules, CA, USA), 1.25 µl of 20,000 ng/µl bovine serum albumin, 0.5 each of 10 µM *rpoB* forward and reverse primers Smyces _rpoB1563F and Smyces_rpoB1968R and 5.75 of molecular grade water. Thermocycler conditions consisted of an initial denaturation step at 98 °C for 30 sec, followed by 40 cycles of 98 °C for 10 sec and 58 °C for 30 sec. Then, a melt curve analysis was performed at increments of 0.5 °C every 5 sec from 65 to 95 °C. Genomic DNA extracted from *Streptomyces scabiei* strain NRRL B-24449 and *Streptomyces lividans* strain NRRL B-65306 (66.1 and 131.146 ng/µl, respectively) was serially diluted between $10^{-1}$ and $10^{-7}$ (1.20 x $10^6$ to 1.2 and 1.31 x $10^6$ and 1.31 gene copies, respectively) and used as standards for all qPCR calculations. The number of *rpoB* gene copies was calculated as previously reported (Ritalahti *et al.*, 2006). The Minimum Information for Publication of Quantitative Real-Time PCR Experiments (MIQE) metrics (Bustin *et al.*, 2009), including slope, squared correlation coefficient, and assay efficiencies, are reported in Table S4 and qPCR standard curves in Fig. S6.

*Pairwise rpoB and MLSA analysis*

Of the total 998 *Streptomyces* genomes downloaded and genes identified in the main text (see main text Materials and Methods), 198 *Streptomyces* genomes were identified as belonging to type strain material in the National Center for Biotechnology Information database and were used in subsequent analyses. A set of five genes (*atpD*, *gyrB*, *recA*, *rpoB*, *trpB*) previously used to discriminate *Streptomyces* species by multi-locus sequence analysis (MLSA) (Rong and Huang, 2012) were downloaded from the PubMLST database (Jolley *et al.*, 2018). The PubMLST gene sequences were aligned to protein-coding genes identified in each *Streptomyces* genome using translated nucleotide alignments with the lastal alignment tool (Kielbasa *et al.*, 2011). Significant alignments (≥50% query amino acid identity, ≥50% query alignment length) were parsed to identify orthologous genes in the 198 *Streptomyces* genomes. We then performed multiple sequence alignment of the full length genomic sequences to their respective PubMLST sequences using MAFFT with default settings (Katoh *et al.*, 2009). Full-length sequences were then trimmed in Jalview (Waterhouse *et al.*, 2009) to encompass the same region of each gene encoded by the PubMLST amplicon sequences. The trimmed sequences for each gene were realigned with MAFFT, aligned regions edited using trimAl (Capella-Gutiérez *et al.*, 2009) with settings '-gt 0.2 -cons 70 -keepheader -keepseqs –phylip', and post-trimAl alignments were concatenated end to end using a custom python script. Pairwise maximum-likelihood evolutionary distance estimates were inferred with the Kimura two parameter (K80) (Kimura, 1980) using RAxML (Stamatakis, 2006) between the concatenated *Streptomyces* MLSA sequences (Rong and Huang, 2012). Pairwise K80 evolutionary distances between *rpoB* nucleotide sequences delimited by PCR primers Smyces _rpoB1563F and Smyces_rpoB1968R were calculated in a similar manner. The *rpoB* pairwise nucleotide identities were regressed onto the pairwise K80 estimates for the full *Streptomyces* MLSA and plotted in R using ggplot2 (Wickham, 2009; R Core Team, 2018) (Fig. S7). An MLSA K80 distance of 0.007 has been previously established as a species cutoff within the genus *Streptomyces* and was shown to equate to a 99% nucleotide identity cutoff for the *rpoB* gene region targeted by the primers Smyces _rpoB1563F and Smyces_rpoB1968R (Fig. S7). All data, including gene sequences, alignments, etc. are available in a figshare repository hosted by S.A.H. (https://doi.org/10.6084/m9.figshare.c.4573250).

### Results and Discussion

*PCR primer design and analysis*

Of the 1,297 *Streptomycetaceae* OTUs identified, ten were classified as the sister genera *Kitasatospora* (n = 5) and *Streptacidiphilus* (n = 5). The number of *Kitasatospora* and *Streptacidiphilus* reads detected were low, and ranged between 6 and 55 (0.001 – 0.013%) and 15 and 346 (0.004 - 0.081%) of total read counts, respectively. Furthermore, we detected *Kitasatospora* and *Streptacidiphilus* OTUs in a total of 27% (n = 4) and 40 % (n = 6) samples, respectively. Although we attempted to maximize the discrimination of *Streptomyces rpoB* sequences over other taxa during *rpoB* primer design (see Materials and Methods main text), a much smaller number of reference *rpoB* sequences were available for members of the

*Kitasatospora* and *Streptacidiphilus*, which likely contributed to our inability to completely exclude these taxa from amplification in the present study.

The three predominant taxa from the non-target fraction were classified as members of the *Pseudonocardiales* (*Actinobacteria*), *Propionibacteriales* (*Actinoacteria*), and *Rhizobiales* (*Alphaproteobacteria*). The *Pseudonocardiales* and *Propionibacteriales* OTUs aligned to *rpoB* genes from representative members of each taxon, but the *Rhizobiales* sequences aligned to *rpoC* gene sequences, a related but distinct subunit of the DNA-directed RNA polymerase. The remaining non-target sequences represented non-specific amplification of DNA sequences other than *rpoB* or *rpoC*. Hence, the amplification of non-target sequences can occur, but at a lower frequency due to nucleotide conservation within the primer design region (Fig. S8). We optimized the annealing temperature in the present study to balance sequence diversity and specificity, but a higher annealing temperature (>55 °C) can be employed to further reduce non-specific amplification with the *rpoB* primer set (or for qPCR). Overall, the contribution of non-target sequences to our dataset was low (< 10%) and the target *Streptomyces rpoB* gene sequences enables robust analyses of *Streptomyces* communities.

### Supplemental Tables

Table S1. Estimated *rpoB* sample coverage and read depth required to reach 95 and 99% sample coverage calculated using the iNEXT tool (Hsieh *et al.*, 2016) for all 15 New Zealand soils.

| Site | Sample coverage (n = 925) | Sample coverage (n = 5000) | Estimated read depth at 0.95 sample coverage | Estimated read depth at 0.99 sample coverage |
|---|---|---|---|---|
| Great Barrier Island 1B | 0.976 | 0.993 | 287 | 3206 |
| Great Barrier Island 4N | 0.978 | 0.993 | 185 | 2936 |
| Oteha Rowe | 0.946 | 0.988 | 1024 | 5732 |
| Ohinewai | 0.946 | 0.984 | 1065 | 8660 |
| Rangipo Desert | 0.986 | 0.997 | 149 | 1373 |
| Mangaweka | 0.978 | 0.995 | 320 | 2253 |
| Pangatotara | 0.968 | 0.99 | 503 | 5217 |
| Hope Saddle Lookout | 0.993 | 0.996 | 152 | 580 |
| Kaikoura Peninsula | 0.957 | 0.993 | 726 | 3986 |
| Woodstock Road | 0.988 | 0.997 | 87 | 1143 |
| Haast Valley | 0.985 | 0.995 | 216 | 1733 |
| Cameron Flats | 0.985 | 0.994 | 163 | 1824 |
| Kawarau Gorge | 0.963 | 0.994 | 640 | 3465 |
| Kingsland Rd | 0.964 | 0.99 | 582 | 4843 |
| Red Tussock Conservation Area | 0.959 | 0.999 | 727 | 2574 |
| ***Mean*** | **0.971** | **0.993** | **455** | **3302** |
| ***Standard deviation*** | **0.015** | **0.004** | **325** | **2127** |
| ***Median*** | **0.976** | **0.994** | **320** | **2936** |
| ***Variance*** | **0.000** | **0.000** | **105610** | **4525772** |
| ***Min*** | **0.946** | **0.984** | **87** | **580** |
| ***Max*** | **0.993** | **0.999** | **1065** | **8660** |

Table S2. The squared correlation coefficients and P-values reported by envfit and adonis permutational tests. Both envfit and adonis functions perform permutations of input ordinations or dissmiliarity matrices, respectively, to assess significance of the correlation of the data with the independent variables. wUniFrac and uUniFrac represent weighted and unweighted UniFrac metrics, respectively.

| **Function** | **Dissimilarity metric** | **Variable** | **$R^2$** | ***P*-value** |
|---|---|---|---|---|
| envfit | Jaccard | pH | 0.49 | **0.016**** |
| | | Temperature | 0.27 | 0.147 |
| | | Rainfall | 0.44 | **0.029**** |
| | | Plant richness | 0.64 | **0.002***** |
| | Bray-Curtis | pH | 0.48 | **0.020**** |
| | | Temperature | 0.32 | 0.102 |
| | | Rainfall | 0.45 | **0.023**** |
| | | Plant richness | 0.65 | **0.003***** |
| | wUniFrac | pH | 0.39 | 0.050* |
| | | Temperature | 0.24 | 0.182 |
| | | Rainfall | 0.75 | **0.002***** |
| | | Plant richness | 0.35 | 0.070* |
| | uUniFrac | pH | 0.50 | **0.014**** |
| | | Temperature | 0.38 | 0.063* |
| | | Rainfall | 0.57 | **0.009***** |
| | | Plant richness | 0.67 | **0.002***** |
| adonis | Jaccard | pH | 0.10 | **0.003***** |
| | | Temperature | 0.07 | 0.162 |
| | | Rainfall | 0.07 | 0.222 |
| | | Plant richness | 0.08 | **0.043**** |
| | Bray-Curtis | pH | 0.11 | **0.003***** |
| | | Temperature | 0.08 | 0.140 |
| | | Rainfall | 0.07 | 0.198 |
| | | Plant richness | 0.09 | 0.048* |
| | wUniFrac | pH | 0.19 | **0.002***** |
| | | Temperature | 0.08 | 0.102 |
| | | Rainfall | 0.20 | **0.003***** |
| | | Plant richness | 0.08 | 0.120 |
| | uUniFrac | pH | 0.13 | **0.002***** |
| | | Temperature | 0.07 | 0.178 |
| | | Rainfall | 0.08 | 0.110 |
| | | Plant richness | 0.08 | 0.119 |

*Note:* $^{*}p<0.1$; $^{**}p<0.05$; $^{***}p<0.01$

Table S3. *Streptomyces rpoB* PCR primers modified for compatibility with Illumina amplicon sequencing as previously described for 16S rRNA gene amplicon sequencing (Kozich *et al.*, 2013).

| Primer label | Kozich barcode label | Sequence (5'-->3') |
|---|---|---|
| rpob.F.SC501 | SC501 | AATGATACGGCGACCACCGAGATCTACACACGACGTGTATGGTAATTGGGGAGGACCGCTTCGTCATC |
| rpob.F.SC502 | SC502 | AATGATACGGCGACCACCGAGATCTACACATATACACTATGGTAATTGGGGAGGACCGCTTCGTCATC |
| rpob.F.SC503 | SC503 | AATGATACGGCGACCACCGAGATCTACACCGTCGCTATATGGTAATTGGGGAGGACCGCTTCGTCATC |
| rpob.F.SC504 | SC504 | AATGATACGGCGACCACCGAGATCTACACCTAGAGCTTATGGTAATTGGGGAGGACCGCTTCGTCATC |
| rpob.F.SC505 | SC505 | AATGATACGGCGACCACCGAGATCTACACGCTCTAGTTATGGTAATTGGGGAGGACCGCTTCGTCATC |
| rpob.F.SC506 | SC506 | AATGATACGGCGACCACCGAGATCTACACGACACTGATATGGTAATTGGGGAGGACCGCTTCGTCATC |
| rpob.F.SC507 | SC507 | AATGATACGGCGACCACCGAGATCTACACTGCGTACGTATGGTAATTGGGGAGGACCGCTTCGTCATC |
| rpob.F.SC508 | SC508 | AATGATACGGCGACCACCGAGATCTACACTAGTGTAGTATGGTAATTGGGGAGGACCGCTTCGTCATC |
| rpob.F.SD501 | SD501 | AATGATACGGCGACCACCGAGATCTACACAAGCAGCATATGGTAATTGGGGAGGACCGCTTCGTCATC |
| rpob.F.SD502 | SD502 | AATGATACGGCGACCACCGAGATCTACACACGCGTGATATGGTAATTGGGGAGGACCGCTTCGTCATC |
| rpob.F.SD503 | SD503 | AATGATACGGCGACCACCGAGATCTACACCGATCTACTATGGTAATTGGGGAGGACCGCTTCGTCATC |
| rpob.F.SD504 | SD504 | AATGATACGGCGACCACCGAGATCTACACTGCGTCACTATGGTAATTGGGGAGGACCGCTTCGTCATC |
| rpob.F.SD505 | SD505 | AATGATACGGCGACCACCGAGATCTACACGTCTAGTGTATGGTAATTGGGGAGGACCGCTTCGTCATC |
| rpob.F.SD506 | SD506 | AATGATACGGCGACCACCGAGATCTACACCTAGTATGTATGGTAATTGGGGAGGACCGCTTCGTCATC |
| rpob.F.SD507 | SD507 | AATGATACGGCGACCACCGAGATCTACACGATAGCGTTATGGTAATTGGGGAGGACCGCTTCGTCATC |
| rpob.F.SD508 | SD508 | AATGATACGGCGACCACCGAGATCTACACTCTACACTTATGGTAATTGGGGAGGACCGCTTCGTCATC |
| rpob.R.SD701 | SD701 | CAAGCAGAAGACGGCATACGAGATACCTAGTAAGTCAGTCAGTAGTACGTGGTGTACGTGCCG |
| rpob.R.SD702 | SD702 | CAAGCAGAAGACGGCATACGAGATACGTACGTAGTCAGTCAGTAGTACGTGGTGTACGTGCCG |
| rpob.R.SD703 | SD703 | CAAGCAGAAGACGGCATACGAGATATATCGCGAGTCAGTCAGTAGTACGTGGTGTACGTGCCG |
| rpob.R.SD704 | SD704 | CAAGCAGAAGACGGCATACGAGATCACGATAGAGTCAGTCAGTAGTACGTGGTGTACGTGCCG |
| rpob.R.SD705 | SD705 | CAAGCAGAAGACGGCATACGAGATCGTATCGCAGTCAGTCAGTAGTACGTGGTGTACGTGCCG |
| rpob.R.SD706 | SD706 | CAAGCAGAAGACGGCATACGAGATCTGCGACTAGTCAGTCAGTAGTACGTGGTGTACGTGCCG |
| rpob.R.SD707 | SD707 | CAAGCAGAAGACGGCATACGAGATGCTGTAACAGTCAGTCAGTAGTACGTGGTGTACGTGCCG |
| rpob.R.SD708 | SD708 | CAAGCAGAAGACGGCATACGAGATGGACGTTAAGTCAGTCAGTAGTACGTGGTGTACGTGCCG |
| rpob.R.SD709 | SD709 | CAAGCAGAAGACGGCATACGAGATGGTCGTAGAGTCAGTCAGTAGTACGTGGTGTACGTGCCG |
| rpob.R.SD710 | SD710 | CAAGCAGAAGACGGCATACGAGATTAAGTCTCAGTCAGTCAGTAGTACGTGGTGTACGTGCCG |
| rpob.R.SD711 | SD711 | CAAGCAGAAGACGGCATACGAGATTACACAGTAGTCAGTCAGTAGTACGTGGTGTACGTGCCG |
| rpob.R.SD712 | SD712 | CAAGCAGAAGACGGCATACGAGATTTGACGCAAGTCAGTCAGTAGTACGTGGTGTACGTGCCG |
| rpob.F.SA501 | SA501 | AATGATACGGCGACCACCGAGATCTACACATCGTACGTATGGTAATTGGGGAGGACCGCTTCGTCATC |
| rpob.F.SA502 | SA502 | AATGATACGGCGACCACCGAGATCTACACACTATCTGTATGGTAATTGGGGAGGACCGCTTCGTCATC |
| rpob.F.SA503 | SA503 | AATGATACGGCGACCACCGAGATCTACACTAGCGAGTTATGGTAATTGGGGAGGACCGCTTCGTCATC |
| rpob.F.SA504 | SA504 | AATGATACGGCGACCACCGAGATCTACACCTGCGTGTTATGGTAATTGGGGAGGACCGCTTCGTCATC |
| rpob.F.SA505 | SA505 | AATGATACGGCGACCACCGAGATCTACACTCATCGAGTATGGTAATTGGGGAGGACCGCTTCGTCATC |
| rpob.F.SA506 | SA506 | AATGATACGGCGACCACCGAGATCTACACCGTGAGTGTATGGTAATTGGGGAGGACCGCTTCGTCATC |
| rpob.F.SA507 | SA507 | AATGATACGGCGACCACCGAGATCTACACGGATATCTTATGGTAATTGGGGAGGACCGCTTCGTCATC |
| rpob.F.SA508 | SA508 | AATGATACGGCGACCACCGAGATCTACACGACACCGTTATGGTAATTGGGGAGGACCGCTTCGTCATC |
| rpob.F.SB501 | SB501 | AATGATACGGCGACCACCGAGATCTACACCTACTATATATGGTAATTGGGGAGGACCGCTTCGTCATC |
| rpob.F.SB502 | SB502 | AATGATACGGCGACCACCGAGATCTACACCGTTACTATATGGTAATTGGGGAGGACCGCTTCGTCATC |
| rpob.F.SB503 | SB503 | AATGATACGGCGACCACCGAGATCTACACAGAGTCACTATGGTAATTGGGGAGGACCGCTTCGTCATC |
| rpob.F.SB504 | SB504 | AATGATACGGCGACCACCGAGATCTACACTACGAGACTATGGTAATTGGGGAGGACCGCTTCGTCATC |
| rpob.F.SB505 | SB505 | AATGATACGGCGACCACCGAGATCTACACACGTCTCGTATGGTAATTGGGGAGGACCGCTTCGTCATC |
| rpob.F.SB506 | SB506 | AATGATACGGCGACCACCGAGATCTACACTCGACGAGTATGGTAATTGGGGAGGACCGCTTCGTCATC |
| rpob.F.SB507 | SB507 | AATGATACGGCGACCACCGAGATCTACACGATCGTGTTATGGTAATTGGGGAGGACCGCTTCGTCATC |
| rpob.F.SB508 | SB508 | AATGATACGGCGACCACCGAGATCTACACGTCAGATATATGGTAATTGGGGAGGACCGCTTCGTCATC |
| rpob.R.SA701 | SA701 | CAAGCAGAAGACGGCATACGAGATAACTCTCGAGTCAGTCAGTAGTACGTGGTGTACGTGCCG |
| rpob.R.SA702 | SA702 | CAAGCAGAAGACGGCATACGAGATACTATGTCAGTCAGTCAGTAGTACGTGGTGTACGTGCCG |
| rpob.R.SA703 | SA703 | CAAGCAGAAGACGGCATACGAGATAGTAGCGTAGTCAGTCAGTAGTACGTGGTGTACGTGCCG |
| rpob.R.SA704 | SA704 | CAAGCAGAAGACGGCATACGAGATCAGTGAGTAGTCAGTCAGTAGTACGTGGTGTACGTGCCG |
| rpob.R.SA705 | SA705 | CAAGCAGAAGACGGCATACGAGATCGTACTCAAGTCAGTCAGTAGTACGTGGTGTACGTGCCG |
| rpob.R.SA706 | SA706 | CAAGCAGAAGACGGCATACGAGATCTACGCAGAGTCAGTCAGTAGTACGTGGTGTACGTGCCG |
| rpob.R.SA707 | SA707 | CAAGCAGAAGACGGCATACGAGATGGAGACTAAGTCAGTCAGTAGTACGTGGTGTACGTGCCG |
| rpob.R.SA708 | SA708 | CAAGCAGAAGACGGCATACGAGATGTCGCTCGAGTCAGTCAGTAGTACGTGGTGTACGTGCCG |
| rpob.R.SA709 | SA709 | CAAGCAGAAGACGGCATACGAGATGTCGTAGTAGTCAGTCAGTAGTACGTGGTGTACGTGCCG |
| rpob.R.SA710 | SA710 | CAAGCAGAAGACGGCATACGAGATTAGCAGACAGTCAGTCAGTAGTACGTGGTGTACGTGCCG |
| rpob.R.SA711 | SA711 | CAAGCAGAAGACGGCATACGAGATTCATAGACAGTCAGTCAGTAGTACGTGGTGTACGTGCCG |
| rpob.R.SA712 | SA712 | CAAGCAGAAGACGGCATACGAGATTCGCTATAAGTCAGTCAGTAGTACGTGGTGTACGTGCCG |
| rpoB_read1_seq_primer | NA | TATGGTAATTGGGGAGGACCGCTTCGTCATC |

| | | |
|---|---|---|
| rpoB_index_seq_primer | NA | CGGCACGTACACCACGTACTACTGACTGACT |
| rpoB_read2_seq_primer | NA | AGTCAGTCAGTAGTACGTGGTGTACGTGCCG |

Table S4. Minimum Information for Publication of Quantitative Real-Time PCR Experiments (MIQE) metrics using the PCR primers Smyces _rpoB1563F and Smyces_rpoB1968R.

| Species | Slope | Y-intercept | $R^2$ | Efficiency |
|---|---|---|---|---|
| *Streptomyces scabiei* strain NRRL B-24449 | -3.34 | 36.3 | 0.9977 | 99.45 |
| *Streptomyces lividans* strain NRRL B-65306 | -3.55 | 35.7 | 0.9997 | 91.3 |

### Supplemental Figures

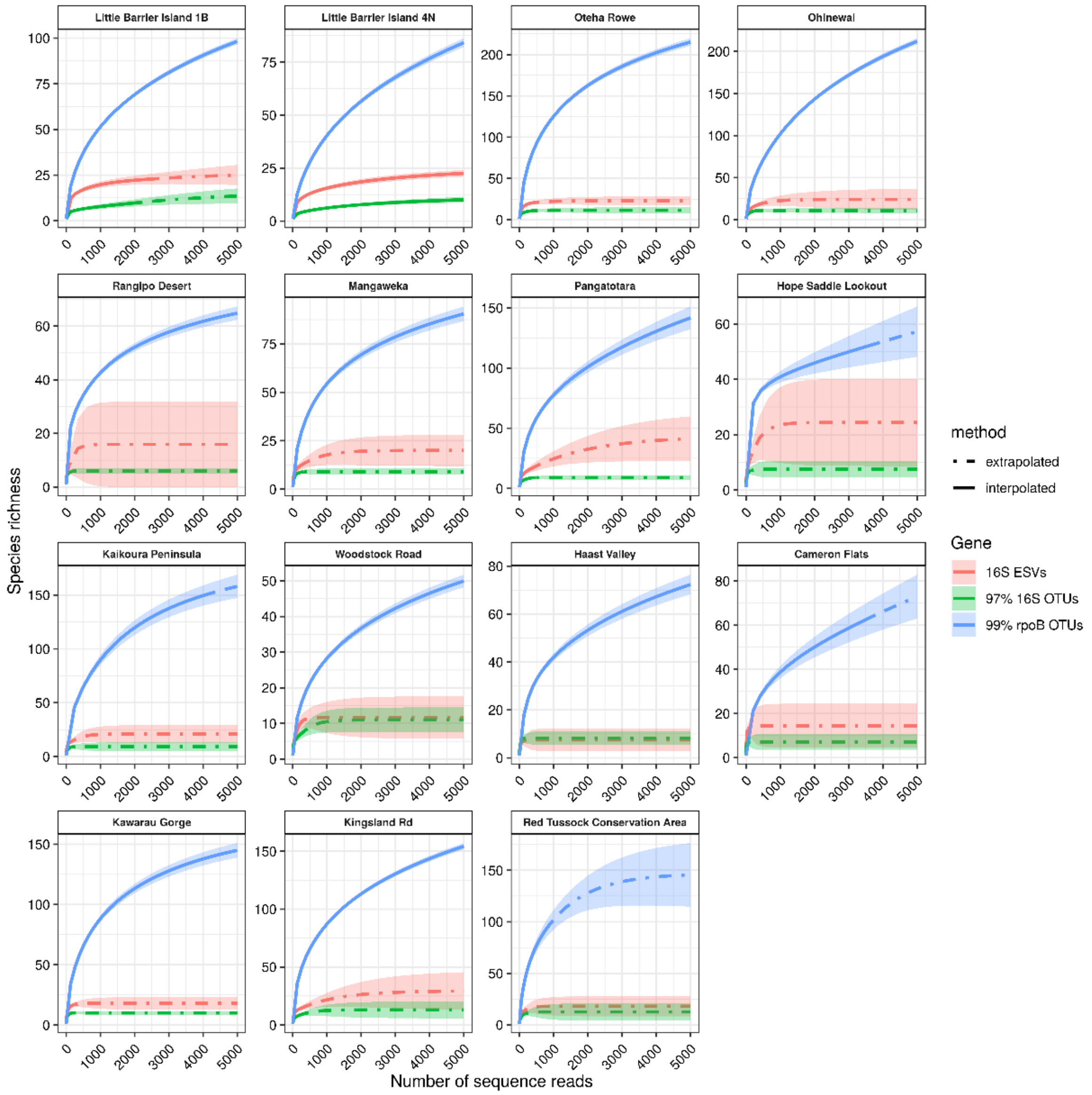


Figure S1. Rarefaction of *Streptomyces rpoB* and 16S gene OTU or exact sequence variant (ESV) richness using the iNEXT tool. The solid and dot-dash lines are interpolated and extrapolated observed richness estimates provided by iNEXT (Chao *et al*., 2014). Since the proportion of *Streptomyces* signal in the 16S gene data sets were low for most samples, we extrapolated beyond limits recommended by iNEXT for qualitative purposes only. The blue, red, and green colors refer to richness estimates for 99% *rpoB* OTUs, 16S gene ESVs, and 97% 16S gene OTUs, respectively.

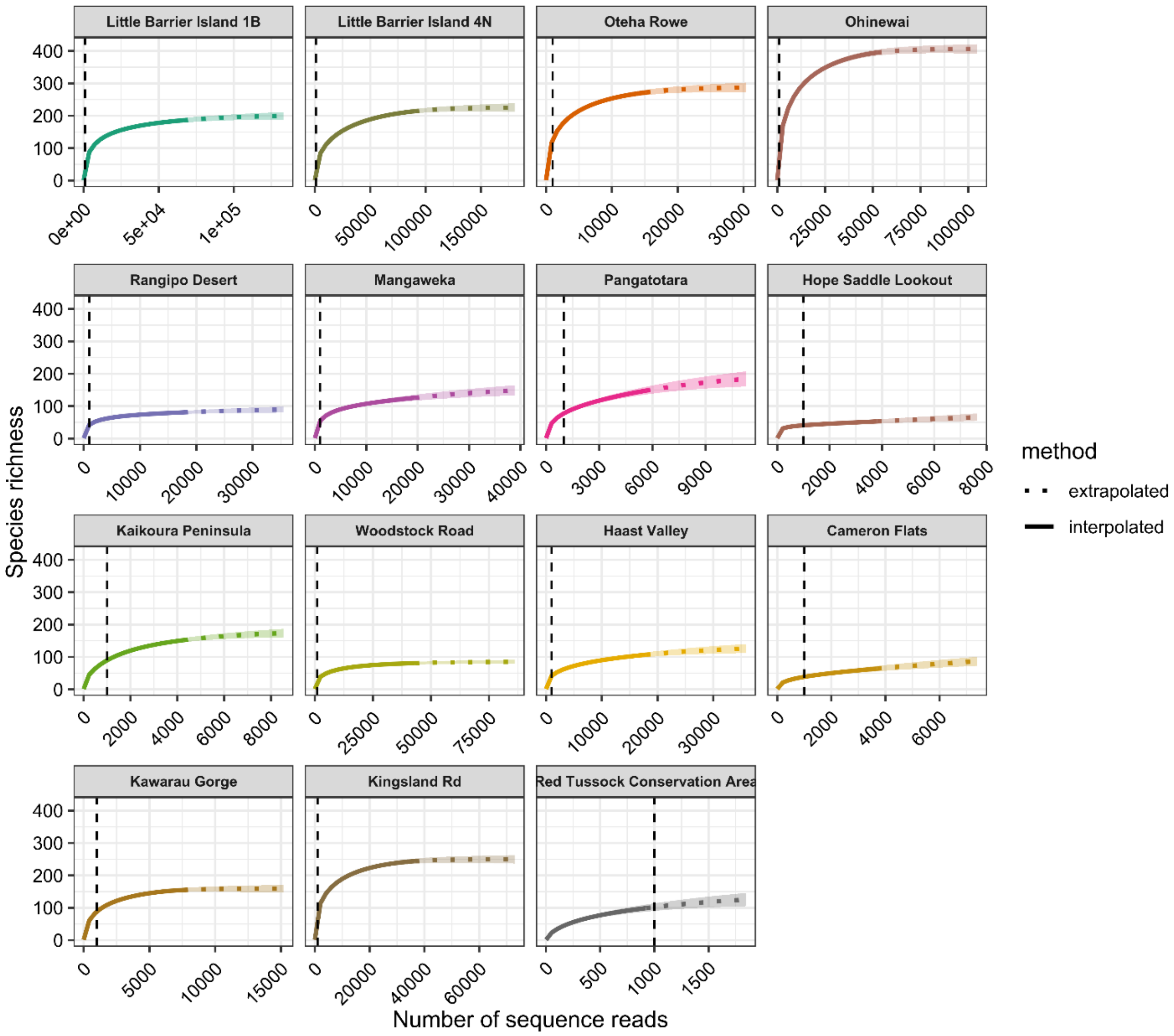


Figure S2. Rarefaction analysis of 99% nucleotide identity clustered *Streptomyces rpoB* OTUs across all 15 New Zealand sites. The solid and dotted lines are interpolated and extrapolated observed richness estimates provided by iNEXT (Chao *et al.*, 2014). Extrapolation was only performed to twice the observed sample size as recommended by iNEXT (Chao *et al.*, 2014). Sites are ordered north to south by latitude starting from the upper left plot to the lower right plot. The vertical dashed line indicates 1,000 sequence reads.

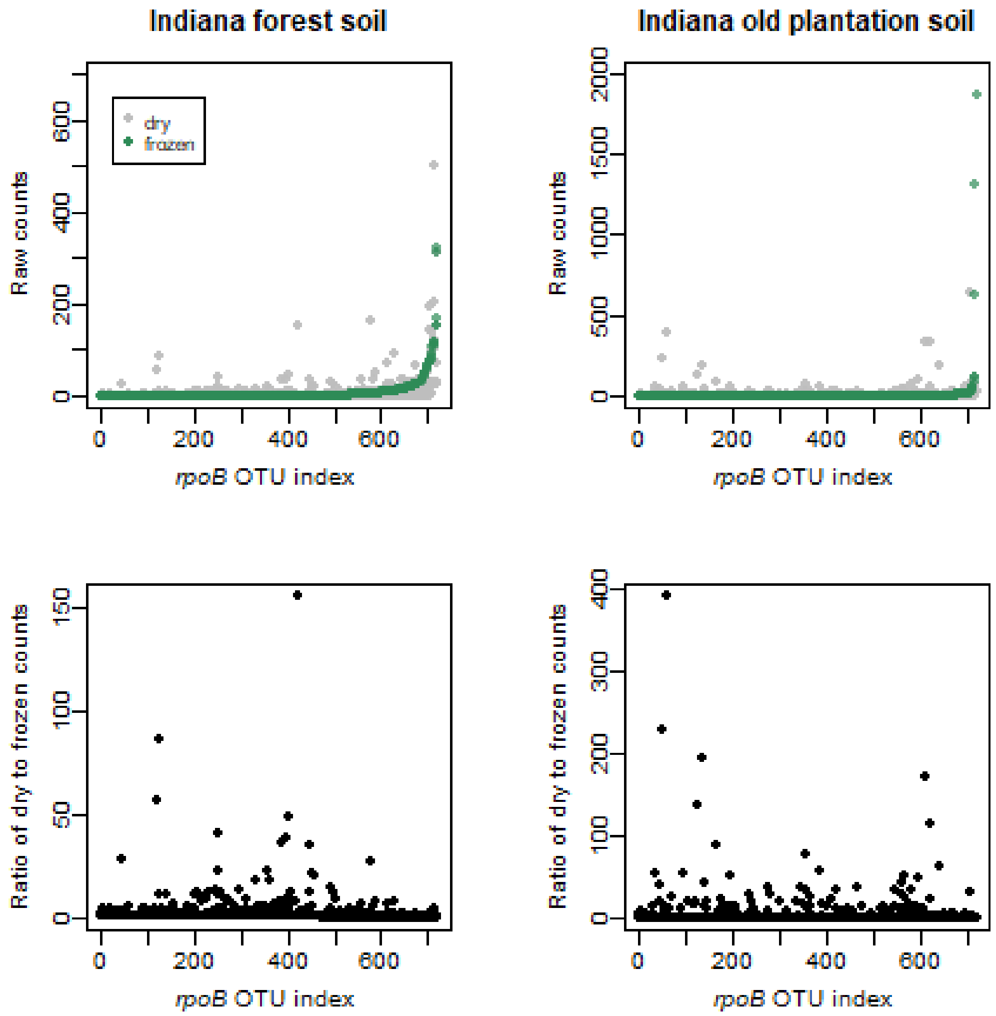


Figure S3. *Streptomyces rpoB* OTU raw reads counts (upper plots) and the ratio of raw read counts (bottom plots) identified within DNA extracted from air-dried or frozen forest (left plots) and plantation soils (right plots), respectively. The *rpoB* OTUs (x-axis) in both samples were sorted by increasing abundance in the frozen samples.

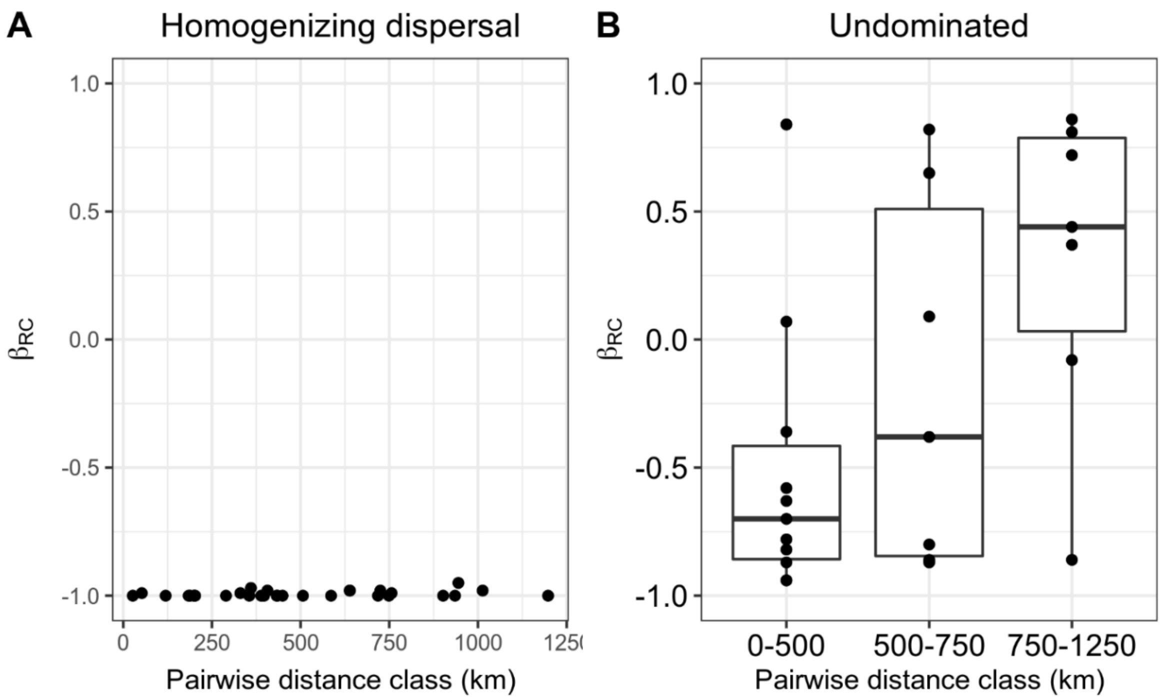


Figure S4. The relationship between pairwise distance and the $\beta_{RC}$ metric for pairwise comparisons classified as "homogenizing dispersal" or "undominated". Briefly, values of $\beta_{RC}$ approaching -1 indicate greater similarity than expected by chance and values approaching 1 indicate much lower similarity than expected by chance. See Stegen et al. (2015) for additional details of the classifications used.

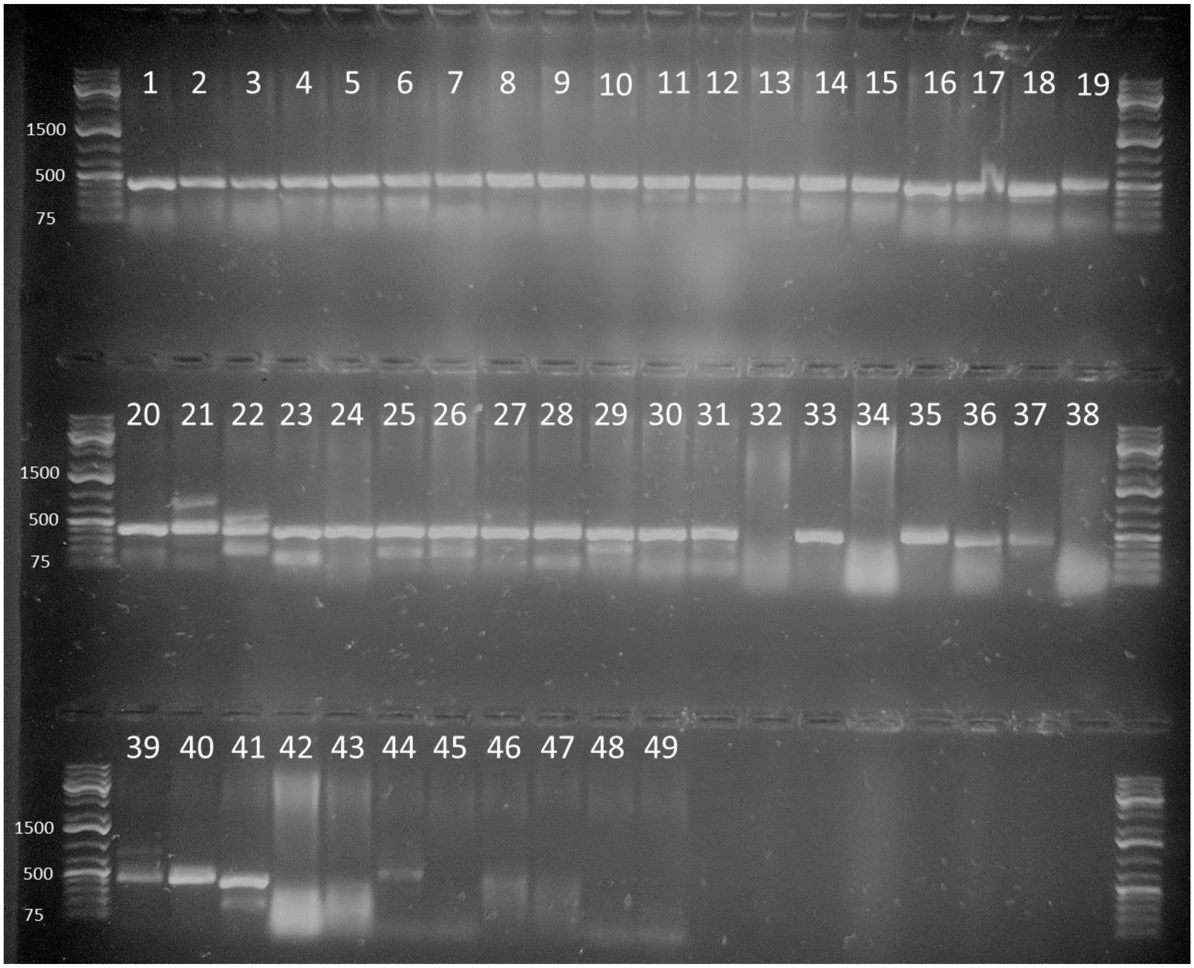


Figure S5. A 1% (w/v) agarose gel containing *rpoB* amplicons from a variety of genomic DNA extracts from bacterial isolates and soil. The values on the left side of the image are molecular marker fragment sizes in nucleotides (GeneRuler 1 KB Plus DNA ladder, ThermoFisher Scientific, Inc). Numbers above wells indicate which genomic DNA sample was loaded into a PCR reaction and their identity is as follows: 1) *Streptomyces* sp. Dul44, 2) *Streptomyces* sp. wa1, 3) *Streptomyces* sp. wa22, 4) *Streptomyces* sp. wa53, 5) *Streptomyces* sp. Adm13, 6) *Streptomyces* sp. ch30, 7) *Streptomyces* sp. Sk2.4, 8) *Streptomyces* sp. man201, 9) *Streptomyces* sp. man147, 10) *Streptomyces* sp. man167, 11) *Streptomyces* sp. ms191, 12) *Streptomyces* sp. sun92, 13) *Streptomyces* sp. sun83, 14) *Streptomyces* sp. sun51, 15) *Streptomyces* sp. ms154, 16) *Streptomyces* sp. me109, 17) *Streptomyces* sp. me47, 18) *Streptomyces* sp. me60, 19) *Streptomyces* sp. or43, 20) *Streptomyces* sp. or59, 21) *Streptomyces* sp. Col6, 22) *Streptomyces* sp. gb1, 23) *Streptomyces* sp. wa51, 24) *Streptomyces* sp. uw30, 25) *Streptomyces* sp. Sk2.1, 26) *Streptomyces* sp. Sk2.17, 27) *Streptomyces* sp. t39, 28) *Streptomyces halstedii* B-1238, 29) *Streptomyces bikiniensis* B-2690, 30) *Streptomyces griseus* B-2682, 31) *Streptomyces flavochromogenes* B-2684, 32) *Kitasatospora arboriphila* NRRL B-24581, 33) *K. mediocidica* NRRL B-16109, 34) *K. griseola* NRRL B-16229, 35) *Streptacidiphilus alcalitolerans* NRRL B-24556, 36) *Streptacidiphilus durhamensis* NRRL B-65271, 37) *Streptacidiphilus griseisporus* NRRL B-24543, 38) *K. cystarginea* NRRL B-16505, 39) *K. azatica* NRRL B-24283, 40)

*Streptacidiphilus oryzae* NRRL B-24636, 41) *Catenulispora acidophila* NRRL B-24433, 42) *Streptacidiphilus albus* NRRL B-24539 , 43) *K. papulosa* NRRL B-16504, 44) C13 soil gDNA , 45) *Aspergillus* sp. gDNA 46) C14 soil gDNA, 47) NxS Microcosm soil gDNA, 48) *Streptococcus mutans* gDNA, 49) No template control.

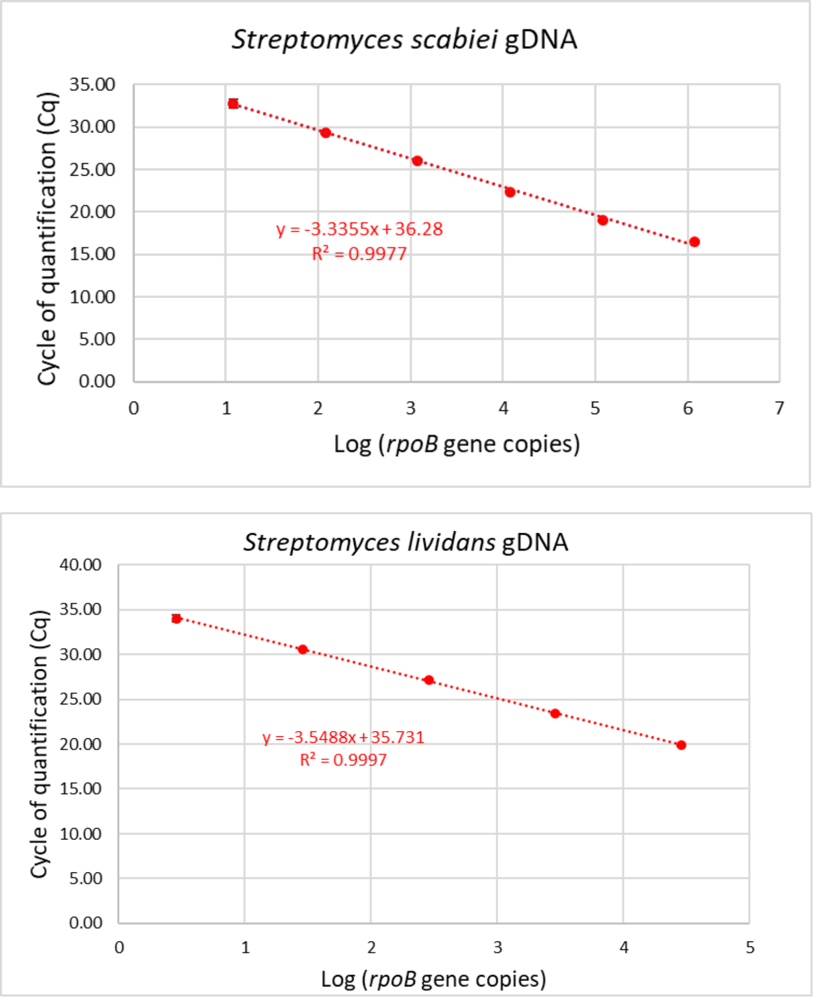


Figure S6. Quantitative PCR standard curves reported for primer pair Smyces _rpoB1563F and Smyces_rpoB1968R for two different *Streptomyces* species. Where visible, black bars above and below points indicate standard deviation from three replicate qPCR reactions.

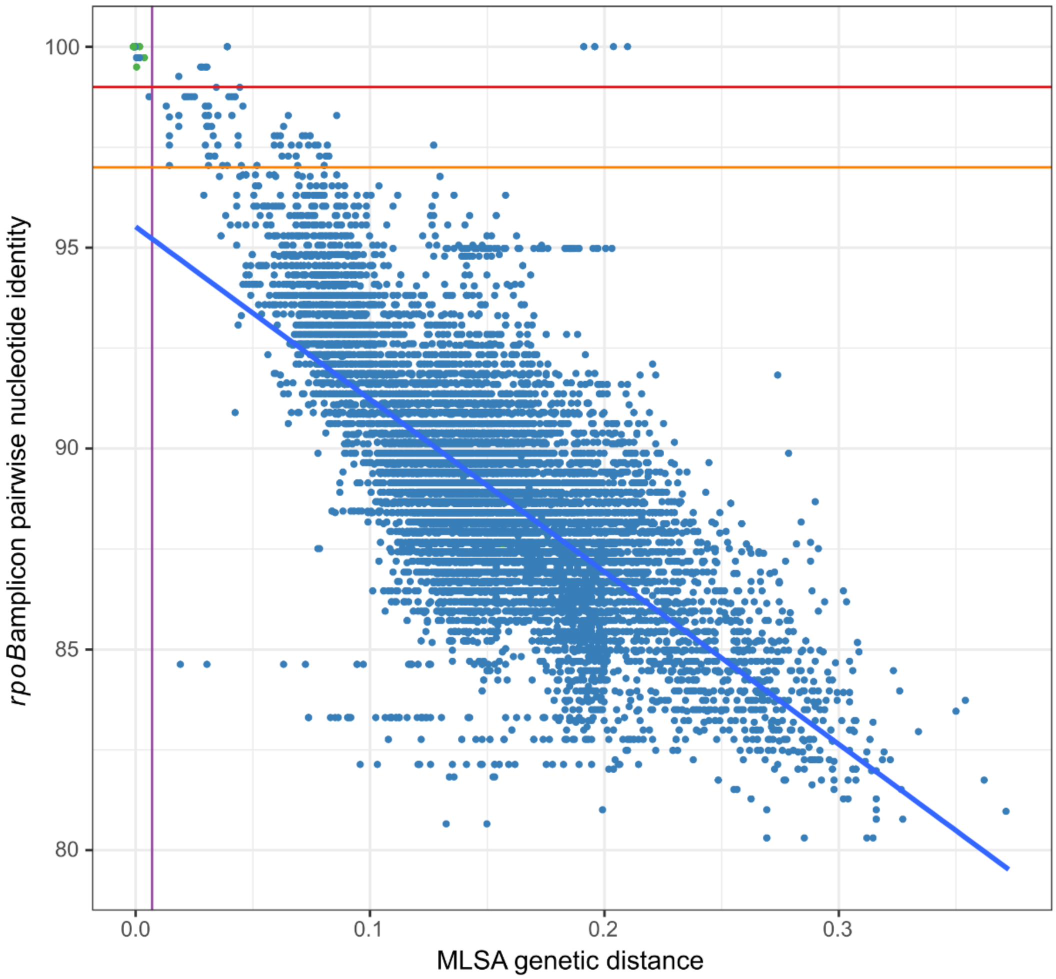


Figure S7. Pairwise nucleotide identity of *Streptomyces rpoB* gene regions targeted by PCR primers Smyces _rpoB1563F and Smyces_rpoB1968R regressed against pairwise K80 distances calculated from a *Streptomyces* MLSA. The solid red and orange horizontal lines indicate a 99% and 97% nucleotide identity cutoff, respectively. The purple vertical line represents a K80 genetic distance of 0.007 previously proposed for species delineation within the *Streptomyces* (Rong and Huang, 2012). The solid blue line represents the best-fit linear regression line. Green and blue dots represent intra- and inter-species comparisons, respectively. Blue dots above the red line and to the right of the purple line may potentially represent species which do not satisfy the 99% nucleotide identity *rpoB* OTU cutoff. It is equally as plausible that these sequences are compromised due to recombination or artifacts due to sequencing and assembly methods. Regardless, few pairwise interactions fall outside of the 99% nucleotide identity and 0.007 genetic distance cutoffs. Blue dots to the left of the purple line and above the red line potentially indicate species of *Streptomyces* that are not presently labeled as conspecifics.

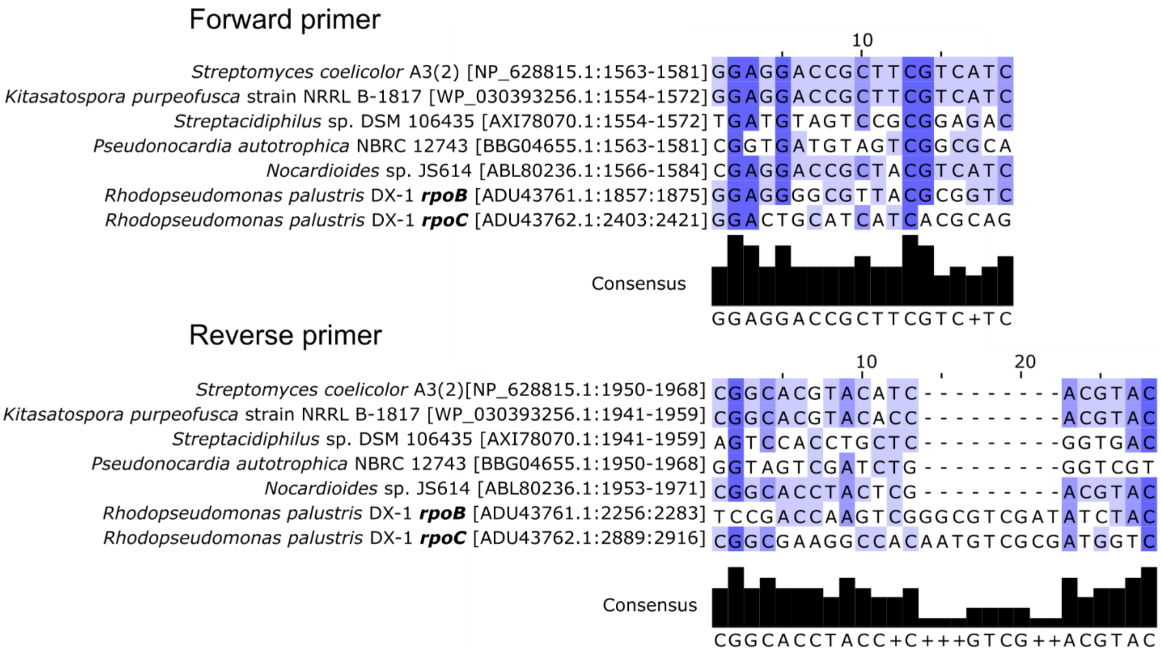


Figure S8. Comparison of *Streptomyces rpoB* PCR primers Smyces _rpoB1563F (5’-GGAGGACCGCTTCGTCATC-3’) and Smyces_rpoB1968R (5’-GTACGTGGTGTACGTGCCG-3’) with representative target (*Streptomyces*) and non-target *rpoB* and *rpoC* nucleotide sequences. The nucleotide sequences in the reverse primer section are the reverse complement of the reverse primer Smyces_rpoB1968R. Values in brackets next to species names indicate the NCBI sequence accession number followed by the alignment position of the primer in the gene’s nucleotide sequence. The *rpoB* and *rpoC* sequences of the alphaproteobacterium *Rhodopseudomonas palustris* strain DX-1 are presented to highlight potential mispriming by sequences of Smyces _rpoB1563F and Smyces_rpoB1968R. Potential mispriming can occur at both 5’ ends of Smyces _rpoB1563F and Smyces_rpoB1968R primers and may indicate why *rpoC* sequences from alphaproteobacterial taxa were identified in the data set. Hence, Illumina sequencing data needs to be filtered to remove any non-target sequences prior to *Streptomyces* community analysis. The remaining filtered sequences can be partitioned by taxonomic classification into *Streptomyces* and other actinobacterial clades amplified by the *rpoB* primer set (e.g., members of the *Kitasatospora* and *Streptacidiphilus*).
